# An intrinsically disordered region of histone demethylase KDM5A activates catalysis through interactions with the nucleosomal acidic patch and DNA

**DOI:** 10.1101/2025.05.01.651538

**Authors:** Ali M Palla, Chien-Chu Lin, Michael J Trnka, Emme M Leao, Nektaria Petronikolou, Alma L Burlingame, Robert K McGinty, Danica Galonić Fujimori

## Abstract

Lysine demethylase 5A (KDM5A) plays a key role in the regulation of chromatin accessibility by catalyzing the removal of trimethyl marks on histone H3K4 (H3K4me3). KDM5A is also an oncogenic driver, with overexpression of KDM5A observed in various cancers, including breast, lung, and ovarian cancer. Past studies have characterized the functions of KDM5A domains, including KDM5A interactions with the histone H3 tail, but have yet to identify the broader mechanisms that drive KDM5A binding to the nucleosome. Through investigation of binding and catalysis on nucleosome substrates, we uncovered multivalent interactions of KDM5A with the H2A/H2B acidic patch and DNA that play crucial roles in the regulation of catalytic activity. We also identified an intrinsically disordered region (IDR) containing bifunctional arginine-rich motifs capable of binding to both the histone H2A/H2B acidic patch and nucleosomal DNA that is necessary for catalysis on nucleosome substrates. Our findings both elucidate previously unknown mechanisms that regulate KDM5A catalytic activity and reveal the ability of an IDR to engage in multiple interactions with chromatin.

**ARTICLE HIGHLIGHTS:**

- The intrinsically disordered region of KDM5A binds the acidic patch and DNA.
- Interactions with the nucleosome are mediated by arginine-rich motifs in the IDR.
- The IDR properly orients KDM5A on the nucleosome to enable catalysis.

## INTRODUCTION

Chromatin structure and accessibility are regulated by histone post-translational modifications (PTMs), which are governed by networks of reader, writer, and eraser proteins. Lysine methylation is one such mark associated with the recruitment of various effectors, leading to transcriptional repression or activation. Removal of lysine methylation is catalysed by lysine demethylase (KDM) proteins, which can be characterized further into families based on their catalytic mechanism, substrate specificity, and domain architecture. The KDM5 family is one such family that removes the trimethylation of lysine 4 on histone H3 (H3K4me3) [1,2]. KDM5 family members (KDM5A-D) share an iron and alpha-ketoglutarate-dependent active site formed by the Jumonji N and C (JmjN and JmjC) domains, a zinc finger domain, a DNA binding AT-rich interaction domain (ARID), and either two or three plant homeodomain (PHD) reader domains (Figure 1A) [3–5]. The ARID and PHD1 domains form a cassette that separates the two Jumonji domains, a feature unique to the KDM5 family [5–7].

**Figure 1:**
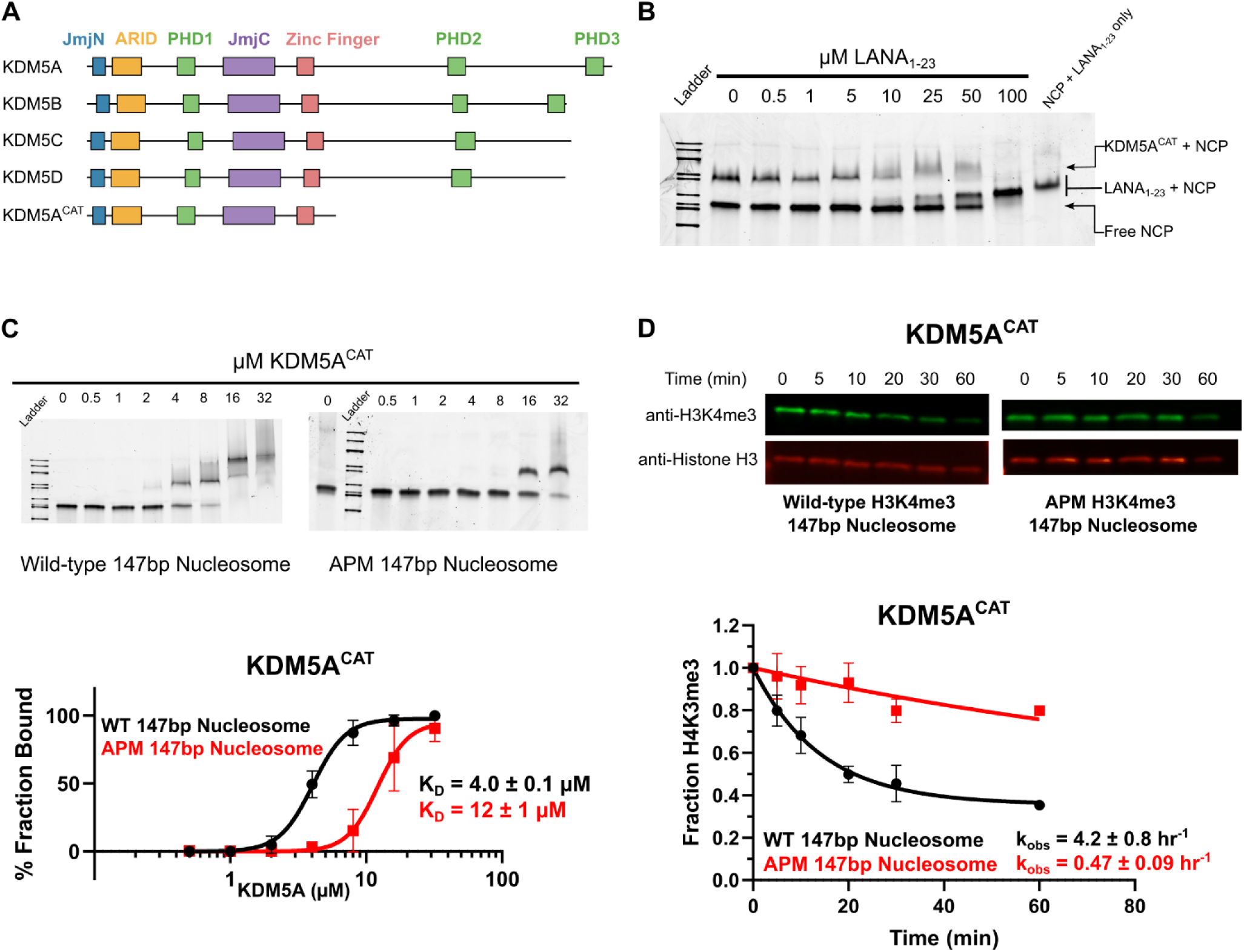
KDM5A interactions with the H2A/H2B acidic patch enhance binding to the nucleosome and stimulate catalytic activity. **A.** Domain architecture of the KDM5 family of proteins and KDM5A^CAT^ construct used for biochemical assays. **B.** EMSA competition assay between KDM5A^CAT^, LANA_1-23_ peptide, and nucleosome. The image is representative of an assay performed in triplicate. Binding reactions were prepared with 8 µM KDM5A and 100 nM wild-type nucleosome, then competed with increasing amounts of LANA peptide. **C.** EMSA binding assay of wild-type KDM5A^CAT^ to wild-type and acidic patch mutant nucleosomes. Data are presented as the mean ± s.d. from four replicates collected across two independent experiments. **D.** KDM5A^CAT^ demethylation of H3K4me3 wild-type and APM 147bp nucleosomes, measured by western blot under single turnover conditions (5 µM KDM5A^CAT^, 50 nM nucleosome substrate, n=2).

KDM5A, in addition to regulating several cellular processes such as differentiation and cell cycle progression, has been linked to various cancer phenotypes [8,9]. In glioma and melanoma cells, KDM5A acts as a tumor suppressor [10,11]. More often, overexpression of KDM5A is an oncogenic driver; high levels of KDM5A are observed across a wide variety of cancerous cells, including glioblastomas, breast, lung, and ovarian cancers [12–17]. A chimeric fusion between the C-terminal PHD3 domain of KDM5A and nucleoporin 98 (NUP98) is frequently observed in acute myeloid leukemia (AML) cells and promotes malignant transformation [18–20]. In glioblastoma and drug-tolerant persister cancer cells, expression of KDM5A confers drug resistance [6,21]. KDM5A thus presents an attractive anti-cancer target due to its role in the progression of a wide variety of cancers [22].

KDM5A contains multiple chromatin binding domains that encompass a variety of functions on nucleosome substrate [4]. The ARID domain displays broad DNA binding ability *in vitro* and recruits KDM5A to certain target genes through recognition of a CG-rich motif [6,21]. Experiments on histone H3K4me3 substrate tail peptides have revealed the substrate preferences of the joint JmjN and JmjC catalytic domain [23–25]. Other domains bind histone H3K4 with alternate functions: PHD1 domain binding allosterically enhances catalytic activity, and the PHD3 domain recruits KDM5A to chromatin [3,26–29]. Despite these findings, the functions of multiple KDM5A domains remain unknown, and a comprehensive portrait of KDM5A regulation on nucleosome substrates has yet to emerge. Moreover, potential interactions of KDM5A with common features targeted by chromatin binding proteins have yet to be investigated, including the H2A/H2B acidic patch, core and flanking DNA, H3 and H2B elbows, and alternate histone tails and modifications [30–32].

Here, by using nucleosomes as physiologically relevant substrates, we uncover previously unobserved interactions of KDM5A and describe how these interactions regulate KDM5A catalytic activity. We find that KDM5A binds to the H2A/H2B acidic patch and extra-nucleosomal linker DNA. We observe that in addition to improving binding affinity, both these features improve the demethylation activity of KDM5A. We also identify bifunctional arginine-rich motifs within an intrinsically disordered region between the ARID and PHD1 domains of KDM5A that interact with the nucleosomal acidic patch and DNA. Combined with previously established interactions, our findings reveal the importance of multivalent interactions that recruit KDM5A to its nucleosome substrate and modulate its catalytic activity.

## MATERIALS AND METHODS

### Cell Culture

*Spodoptera frugiperda* (Sf21) cells (Expression Systems 94-003S) were cultured in ESF921 media with no additives (Expression Systems 96-001) in suspension at 27°C, rotating at 125 rpm.

### Baculoviral expression and purification of recombinant KDM5A constructs

All KDM5A catalytic constructs were expressed in Sf21 cells using a protocol adapted from the Gibco Bac-to-Bac Baculovirus Expression system and previously described [33]. KDM5A sequences were cloned using NEB HiFi DNA Assembly (NEB E2621) into an empty pFastBac vector (pFB-LIC-Bse, Opher Gileadi, Addgene plasmid # 26108) containing a 6xHis tag and TEV cleavage site. The created pFastBac-KDM5A plasmid was transformed into DH10Bac Competent Cells (Gibco 10361012). Bacmid was then purified from DH10Bac cells using Zymo ZR Bac Miniprep Kit (Zymo D4048), followed by transfection into Sf21 cells. For transfection, approximately 8 × 10^5^ Sf21 cells were plated in 2.5 mL of ESF921 media. 8 μL of Cellfectin II reagent (Gibco 10362100) in 100 μL ESF921 was mixed with 3 μg of bacmid in 100 μL ESF921 and incubated at room temperature for 15-30 minutes. Bacmid-Cellfectin mixture was added dropwise to the plated cells, after which cells were incubated at 27°C for 5 hours. All media was then removed from the cells and replaced with fresh ESF921. Transfected cells were incubated for either 5 days or until signs of infection were visible, after which the cells were harvested and spun down to pellet cells. The supernatant was sterile filtered to generate P1 viral stock. P2 viral stock was generated by infecting 25-50 mL of Sf21 suspension culture at 2 × 10^6^ cells/mL with P1 virus at a 20 mL/L ratio of virus:culture. Virus was then harvested after two days, or when cells showed signs of infection, by sterile filtering the supernatant. P3 viral stock was generated in a similar manner, except through infection of 100-200 mL of Sf21 suspension culture using P2 virus. Finally, 800 mL-1 L of cells were infected with P3 virus at a 40mL/L virus:culture ratio for 48 hours. Cells were collected via centrifugation at 1000 rpm, washed with 1x PBS, pelleted again, flash frozen in liquid nitrogen, and stored at -80°C until purification.

For purification, frozen cells were thawed and resuspended in lysis buffer (25 mM HEPES pH 7.9, 350 mM NaCl, 5 mM KCl, 1.5 mM MgCl_2_, 5 mM imidazole, Pierce™ Protease Inhibitor Tablets, EDTA-free). Benzonase (MilliporeSigma 70-746-3) was added to a final concentration of 0.025 U/mL, after which cells were passed through a 70 μm nylon cell strainer (Falcon 352350). Cells were then lysed by performing three passes at 3000 psi through an LM10 Microfluidizer (Microfluidics), followed by centrifugation at 140,000g and filtration through a 0.45 μm PES filter to clarify the lysate. Clarified lysate was incubated with cobalt resin (TALON® Metal Affinity Resin, Takara Bio, 635503) equilibrated in lysis buffer for 1hr at 4°C, and His-tagged KDM5A was eluted with elution buffer (25 mM HEPES pH 7.9, 100 mM NaCl, 0.5 mM MgCl_2_, 10% glycerol, 0.5 mM TCEP) in a stepwise gradient of imidazole (20mM to 200mM Imidazole). His-tag was cleaved through overnight incubation with TEV protease at 4°C in dialysis buffer (25 mM HEPES pH 7.9, 200 mM NaCl, 10% glycerol, 0.5 mM TCEP). The digestion product was passed through cobalt resin, and the flow-through containing cleaved KDM5A enzyme was collected. Protein was purified further by size-exclusion chromatography (HiLoad 16/600 Superdex 200, Cytiva) in storage buffer (2 mM HEPES pH 7.5, 50 mM KCl). Fractions of the highest purity were pooled and concentrated in Amicon Ultra Centrifugal Filters, with a 50 kDa molecular weight cut off, flash-frozen in liquid nitrogen, and stored at -80°C.

pFB-LIC-Bse was a gift from Opher Gileadi (Addgene plasmid # 26108).

### Expression and purification of F40 sortase

F40 sortase was purified as previously described [34]. Briefly, F40 sortase A was overexpressed in BL21 *E. coli*, followed by lysis in 20 mM Tris pH 8.0, 0.1% Triton X-100, and Roche protease inhibitor cocktail. Cleared lysate was applied to high-binding-capacity IMAC resin (Ni Sepharose 6 Fast Flow, Cytiva), washed with 20 mM Tris-HCl pH 7.5, 500 mM NaCl, and eluted in 20 mM Tris-HCl pH 7.5, 150 mM NaCl with increasing amounts of imidazole (up to 250 mM). Fractions containing F40 sortase were combined, dialyzed against 50 mM Tris-HCl pH 7.5, 150 mM NaCl, 5 mM CaCl_2_, concentrated, and stored at -80°C until use.

### Semi-synthesis of H3K4me3 histone

H3K4me3 histones were generated through a semi-synthesis of methylated Histone H3 depsipeptides (aa 1-34) and globular H3 fragment (aa 33-135) by F40 sortase. Histone H3 depsipeptide was purchased from Genscript, comprising the first 34 residues of Histone H3 with the substitution of the amide bond between T33 and G34 with an ester and C-terminal amidation. Globular H3 fragment (gH3) was expressed and purified following established protocols [35].

H3K4me3 histones were synthesized using a previously described protocol with modifications [34]. Histone H3 depsipeptide was diluted into 50 mM HEPES, pH 7.5 and 1 mM DTT, followed by subsequent addition of gH3 to a final concentration of 70 μM, NaCl to 150 mM, and CaCl_2_ to 5 mM. The reaction was initiated by the addition of F40 sortase to a final concentration of 300μM.

The reaction mixture was incubated at 37°C overnight. The following morning, the precipitate was pelleted and resuspended in IEX Buffer (20 mM Tris, pH 7.8, 1 mM EDTA, 7M Urea, 5 mM BME) containing 100 mM NaCl. The sample was loaded onto a cation exchange chromatography column (HiTrap SP HP 5 mL, Cytiva) and eluted using a gradient of IEX buffer containing 100-500 mM NaCl. Fractions containing ligated H3 were pooled, dialyzed against water, and lyophilized before storage at -80°C.

### Nucleosome assembly

Recombinant human 5’ biotinylated 147bp unmodified mononucleosomes (16-0006), 5’ biotinylated 147bp H3K4me3 mononucleosomes (16-0316), and 5’ biotinylated 187bp H3K4me3 mononucleosomes (Custom Order 20-4000) were purchased from Epicypher. Non-biotinylated unmodified and H3K4me3 human nucleosomes were assembled using 147bp or 185bp Widom 601 sequence, containing 1 or 20bp symmetric DNA linkers, respectively, and recombinant or semisynthetic human histones (H2A or H2A APM, H2B, H3.2 or H3K4me3, and H4) as previously described [36]. Briefly, wild-type, unmodified histones and the H2A acidic patch mutant (APM = E61A, E64S, N68A, D72S, N89A, D90A, E91S) histone were expressed in *Escherichia coli* BL21(DE3)pLysS cells at 37°C for 3 h and extracted from washed inclusion bodies as previously described [37]. H2A-H2B (or H2A APM-H2B) dimers and H3-H4 tetramers were reconstituted by dialysis into refolding buffer (10 mM HEPES, pH 7.5, 100 mM NaCl, 10 mM 2-mercaptoethanol) and purified using cation-exchange chromatography (Source S resin, GE Healthcare, 17094405). Nucleosome reconstitution was performed using salt gradient dialysis with purified H2A-H2B dimers and H3-H4 tetramers and either 147bp or 185bp DNA. Reconstituted nucleosomes were subsequently purified using anion-exchange chromatography (Source Q resin, GE Healthcare, 17094705). Pooled fractions were dialyzed into nucleosome storage buffer (10 mM potassium cacodylate, pH 6.5, 0.1 mM EDTA), concentrated to ∼10mg/mL, and stored following the addition of glycerol to a 20% final concentration.

### Electrophoretic mobility shift assays (EMSAs)

For competition binding assays, 100 nM nucleosome, 8 µM KDM5A^CAT^, and varying concentrations of LANA_1-23_ peptide were prepared in binding buffer (50 mM HEPES pH 7.5, 50 mM KCl, 1 mM BME, 0.01% Tween-20, 0.01% BSA, 5% sucrose). For direct nucleosome binding, 100 nM nucleosome and varying concentrations of KDM5A were combined in binding buffer. Both reactions were incubated for 1 hour on ice, then separated on a 7.5% precast protein gel (Mini-PROTEAN TGX, Bio-Rad) at 100V for 2 hours at 4°C in 0.5x Tris-Glycine buffer. For DNA binding, 100 nM DNA and varying concentrations of KDM5A were incubated in TE buffer with 10% glycerol, then run on a 4-20% precast protein gel (Mini-PROTEAN TGX, Bio-rad) at 100V for 80 minutes at 4°C in 0.5x Tris-Glycine buffer. Prior to the addition of the sample, all gels were pre-run at 100V for 30 minutes at 4°C. Following electrophoresis, gels were stained with SYBR gold and imaged using a Chemidoc imaging system, and bands were quantified using ImageLab. To determine IC_50_ in competition binding assays, the fraction of KDM5A bound to nucleosome was fitted to the “One site - logIC50” model in GraphPad Prism. For dissociation constants from direct binding assays, the fraction of bound nucleosome or DNA was fitted to the “Specific binding with hill slope” model in GraphPad Prism.

### Lysine cross-linking and mass spectrometry

Thirty micrograms of KDM5A^CAT^ were crosslinked with a 50-fold molar excess of BS^3^ (ThermoFisher A39266) for 2 hours at 4°C with and without a 10-fold molar excess of ARID-C1 DNA. Crosslinking reactions were quenched by adding Tris-HCl pH 7.5 to a final concentration of 50 mM, followed by precipitation in -20°C acetone. Crosslinked protein was pelleted by TBST centrifugation at 15,000g, washed once with cold acetone, dried, and resuspended in 6 M Guanidine-HCl, 100 mM Ammonium Bicarbonate (ABC), and 10 mM DTT. The solution was then incubated at 56°C for 20 minutes, alkylated in 15 mM iodoacetamide for 45 minutes, and quenched in 25 mM DTT. The sample was diluted to 2 M Guanidine-HCl using ABC and digested by the addition of 0.6 micrograms of LysC, followed by incubation for 4 hours at 37°C. The digestion reaction was diluted again to <1 M Guanidine-HCl using ABC, and 1 microgram of Trypsin was added, followed by incubation at 37°C overnight. The digested sample was acidified to a pH ≤ 3 using formic acid, desalted using C18 stage tips (C18 Empore 6091), and evaporated to dryness. Dried protein was resuspended in 250 mM HEPES, pH 8.5, and labelled with 120 micrograms of Tandem Mass Tag (TMT) 6-plex reagents (Thermo 90061) for 1 hour at room temperature. TMT labeling reactions were quenched by the addition of Tris-base to 100 mM final. Labelled proteins were then acidified to a pH ≤ 3 using formic acid, desalted using C18 stage tips (Empore 6091), evaporated to dryness, and resuspended in 10 μL 0.1% formic acid.

Samples were resuspended in 0.1% formic acid for injection into an Orbitrap Exploris 480 mass spectrometer (Thermo) coupled through an EASY-Spray nano ion source (Thermo) and a FAIMS source (Thermo) to a Dionex UltiMate 3000 uPLC (Thermo) running an EASY-Spray column (75 µm x 50 cm column packed with 2 µm, 100 Å PepMap C18 resin; Thermo). The mobile phases were: solvent A: water/0.1% formic acid; solvent B: acetonitrile/0.1% formic acid. Each sample was loaded at 300 nL/min at 2% B, and then eluted with a 200 nL/min gradient from 2-35% B over 180 min. The column was then washed at 85% B and re-equilibrated back to 2%. The total run time was 237 minutes. Precursor ions were acquired from 350-1500 m/z in the Orbitrap (120k resolving power, 100% AGC, 50ms max injection time). The FAIMS source was operated at 3 CVs (-45, -60, -75 V) with product ions acquired for 1 sec cycle times at each CV. Precursor ions with charges 3-8+ and intensity greater than 50,000 were isolated in the quadrupole (1.6 m/z selection window) and dissociated by HCD with stepped 23, 30, 45% NCE. Product ions were acquired in the Orbitrap (30k resolving power, 100% AGC, 150 ms max injection time). A 30s dynamic exclusion window and the peptide monoisotopic ion precursor selection option were enabled.

Peaklists in MGF format were generated by PAVA with the option to recalculate monoisotopic peak assignments using Monocle [38]. A restricted database consisting of the 20 most abundant proteins identified in the sample was searched for BS3 crosslinks. A decoy database consisting of 20 randomized sequences (which were each 10x longer than their corresponding target protein sequences) was used to model the distribution of incorrect hits and assess FDR. Peaklists were searched with Protein Prospector v6.5.2 with Trypsin specificity and 2 missed cleavages. Precursor and product ion tolerances were 10 and 25 ppm, respectively. DSS crosslinking was specified with 1,000 intermediate hits saved. Carbamidomethylation of Cys was specified as a constant modification. Variable modifications were: Met oxidation, loss and/or acetylation of protein N-terminal Met, peptide N-terminal Glu conversion to pyroglutamate, dead-end DSS modification at Lys and protein N-terminus, incorrect monoisotopic peak assignment (neutral loss of 1Da), and modification by the TMT-6plex reagent (at Lys and the peptide N-terminus). Up to 6 variable modifications per peptide were allowed. Crosslinked matches were classified using Touchstone (in-house R package, manuscript in development) with a minimum peptide length of 3 amino acids and a minimum number of 3 backbone bond cleavages observed per peptide. Crosslinks were summarized at the unique-residue pair level and reported at 1% FDR. The TMT reporter ion intensities were extracted from the peaklists using an in-house Python script. Log2 transformed intensities were median normalized for each channel and the fold change was reported as the average of the Log2 transformed difference between the +DNA channel and a matched -DNA channel for three biological replicates. Only crosslinks discovered in all three experiments were included.

### Demethylation of H3 peptide substrate

Demethylation kinetics were measured using a formaldehyde dehydrogenase assay, coupling the demethylation of H3 peptide substrate to the formation of formaldehyde as described previously [23]. 1 μM KDM5A catalytic construct and 100 μM H3K4me3 peptide (aa 1-21, Genscript) were combined in a demethylation buffer containing 50 mM HEPES pH 7.5, 50 mM KCl, 50 μM (NH_4_)_2_Fe(SO_4_)_2_, 1 mM α-ketoglutarate, 2 mM ascorbic acid, 2 mM NAD+ and 0.05 U formaldehyde dehydrogenase (Sigma-Aldrich F1879). Reactions were initiated by the addition of an enzyme cocktail ((NH_4_)_2_Fe(SO_4_)_2_, ascorbic acid, NAD+, formaldehyde dehydrogenase, and KDM5A) to the substrate mix (peptide and α-ketoglutarate). Formation of formaldehyde was measured by monitoring fluorescence on a SpectraMax M5e (Molecular Devices) using 350 nm excitation and 460 nm emission wavelengths in 20-second intervals. Reaction mixtures without peptide were used as a negative control and for baseline correction, and all conditions were performed in triplicate.

### Demethylation of nucleosome substrate measured by western blot

Demethylation reactions on nucleosome substrate were performed in 50 mM HEPES pH 7.5, 50 mM KCl, 100 μM (NH_4_)_2_Fe(SO_4_)_2_, 100 μM α-ketoglutarate, and 1 mM ascorbic acid. Reactions under single turnover conditions were performed on 50 nM H3K4me3 nucleosomes and 5 μM KDM5A. Demethylation was initiated by the addition of enzyme/cofactor master mix ((NH_4_)_2_Fe(SO_4_)_2_, α-ketoglutarate, ascorbic acid, and KDM5A) to nucleosome substrate and quenched by the addition of 6x SDS buffer.

Reactions were analysed by western blot on 0.2 μm nitrocellulose membranes. Membranes were incubated in primary antibodies anti-H3K4me3 (1:2000, Abcam ab8580) and anti-Histone H3 (1B1B2) (1:1000, CST 14269), overnight at 4°C in 5% Dry Milk (Bio-rad #1706404), followed by incubation in secondary antibodies anti-Rabbit AlexaFluor Plus 800 (1:5000, ThermoFisher A32735) and anti-Mouse AlexaFluor Plus 647 (ThermoFisher A32728) for 1 hour at RT in TBST. Membranes were then imaged on a Chemidoc imaging system (Bio-rad), and signal intensity was quantified using ImageLab. Intensity of H3K4me3 signal was normalized to H3 signal, then again normalized to the 0-time point; values are reported as a relative fraction of H3K4me3 relative to t=0. For experiments where multiple time points were collected, rates were determined by fitting the data to a “One-phase decay” model in GraphPad Prism.

### Demethylation of nucleosome substrate measured by TR-FRET

Demethylation reactions on nucleosome substrate were performed in 50 mM HEPES pH 7.5, 50 mM KCl, 0.01% Tween-20, 0.01% BSA, 100 μM (NH_4_)_2_Fe(SO_4_)_2_, 100 μM α-KG, and 1 mM ascorbic acid. Various concentrations of KDM5A were reacted with 25 nM of biotinylated nucleosome substrate; reactions were initiated by the addition of the enzyme. All reactions were carried out under single-turnover conditions (enzyme in at least 10-fold excess of substrate). Time points were collected by quenching 5 μL of reaction mixture in 1.3 mM EDTA, followed by dilution to 20 μL in 0.5X LANCE detection buffer (PerkinElmer CR97) and the addition of detection reagents 1 nM LANCE Ultra Europium anti-H3K4me1/2 antibody (PerkinElmer TRF0402) and 50 nM LANCE Ultra Ulight-Streptavidin (PerkinElmer TRF0102). Reaction mixtures with detection reagents were transferred to a 384-well white microplate (PerkinElmer 6007290) and incubated in the dark at RT with detection reagents for 2 hours prior to TR-FRET detection. TR-FRET emission was measured at 665 nm and 615 nm with 320 nm excitation, 50 µs delay, and 100 µs integration time, using a Molecular Devices SpectraMax M5e plate reader. FRET ratios were calculated as the 665/615 emission ratio, initial rates were determined using the first 5 minutes, then fit into an “Allosteric sigmoidal kinetics” model in GraphPad Prism.

### AlphaFold3 structure generation and *in silico* screen of acidic patch interactors

AlphaFold3 was run on the AlphaFoldServer (Google) [39]. Protein sequences used are recorded in Supplementary Table S1. Acidic patch interacting residues of the aforementioned structures were screened for using a geometric triangle analysis described previously [40].

## RESULTS

### The H2A/H2B acidic patch stimulates KDM5A catalysis

We began investigating KDM5A interactions with the nucleosome by assaying for binding between KDM5A and the H2A/H2B acidic patch, a frequent interactor with chromatin-modifying proteins. To probe potential KDM5A binding to the acidic patch, we evaluated the ability of LANA_1-23_ peptide, a previously characterized acidic patch interactor [41], to displace interactions between the nucleosome core particle (147bp nucleosome) and KDM5A in an electrophoretic mobility shift assay (EMSA) (Supplementary Figure S1A). In these experiments, we utilized a minimal catalytically active KDM5A 1-801 construct, hereon referred to as KDM5A^CAT^, consisting of the catalytic JmjN and JmjC domains and the ARID, PHD1, and zinc finger domains (Figure 1A). LANA_1-23_ disrupted KDM5A binding to 147bp nucleosome in a dose-dependent manner (Figure 1B). Assays performed with an acidic patch binding-deficient peptide, LANA_1-23_ R9A, demonstrated an inability of the mutant peptide to displace KDM5A interactions with the nucleosome (Supplementary Figure S1B). We then reconstituted acidic patch mutant nucleosomes (APM Nucleosomes: H2A E61A, E64S, N68A, D72S, N89A, D90A, E91S) and assayed binding to KDM5A, uncovering an approximately 3-fold loss in affinity in APM 147bp nucleosomes relative to wild-type, further confirming KDM5A binding to the H2A/H2B acidic patch (Figure 1C). Binding of KDM5A^CAT^ to wild-type H3K4me3 nucleosomes displayed weaker affinity relative to the corresponding unmodified nucleosomes (Supplementary Figure S2). Importantly, similarly to unmodified nucleosomes, mutation of the acidic patch results in a 2-fold lower affinity of KDM5A^CAT^ for APM H3K4me3 modified vs wild-type H3K4me3 nucleosomes.

The observation that KDM5A interacts with the acidic patch prompted us to evaluate the impact of this interaction on the catalytic activity of the demethylase. Demethylation was quantified through quantitative western blot assays with antibodies targeting H3K4me3 and Histone H3, using amounts of nucleosome within the linear range of detection for both antibodies (Supplementary Figure S3). Under single turnover conditions (5 μM KDM5A enzyme, 50 nM substrate), we observed significantly decreased demethylation of H3K4me3 on APM 147bp nucleosomes relative to the corresponding wild-type nucleosomes. The observed approximate 9-fold reduction indicates that the acidic patch stimulates catalytic activity of the enzyme (Figure 1D).

### An intrinsically disordered region between the ARID and PHD1 domains binds to the H2A/H2B acidic patch and regulates catalytic activity

Acidic patch interactions are mediated by motifs including nearby arginine residues, with a canonical anchor in a P0 position and at least one variant arginine, often in a P ± 2 or 3 position [30,31]. We identified three instances of such a motif in an interdomain region between the ARID and PHD1 domains. Despite the overall low conservation of this region across species, these arginine residues are highly conserved in KDM5A orthologs across vertebrates and partially conserved among KDM5 family members in humans (Figure 2A, Supplementary Figure S4). KDM5 family members have been crystallized previously, however, the region between the ARID and PHD1 domain remains unresolved [2,42–44]. Disorder predictions reveal the region between the ARID and PHD1 domains is an intrinsically disordered region (IDR), and this IDR is present in all four human paralogs of the KDM5 family (Figure 2B).

**Figure 2:**
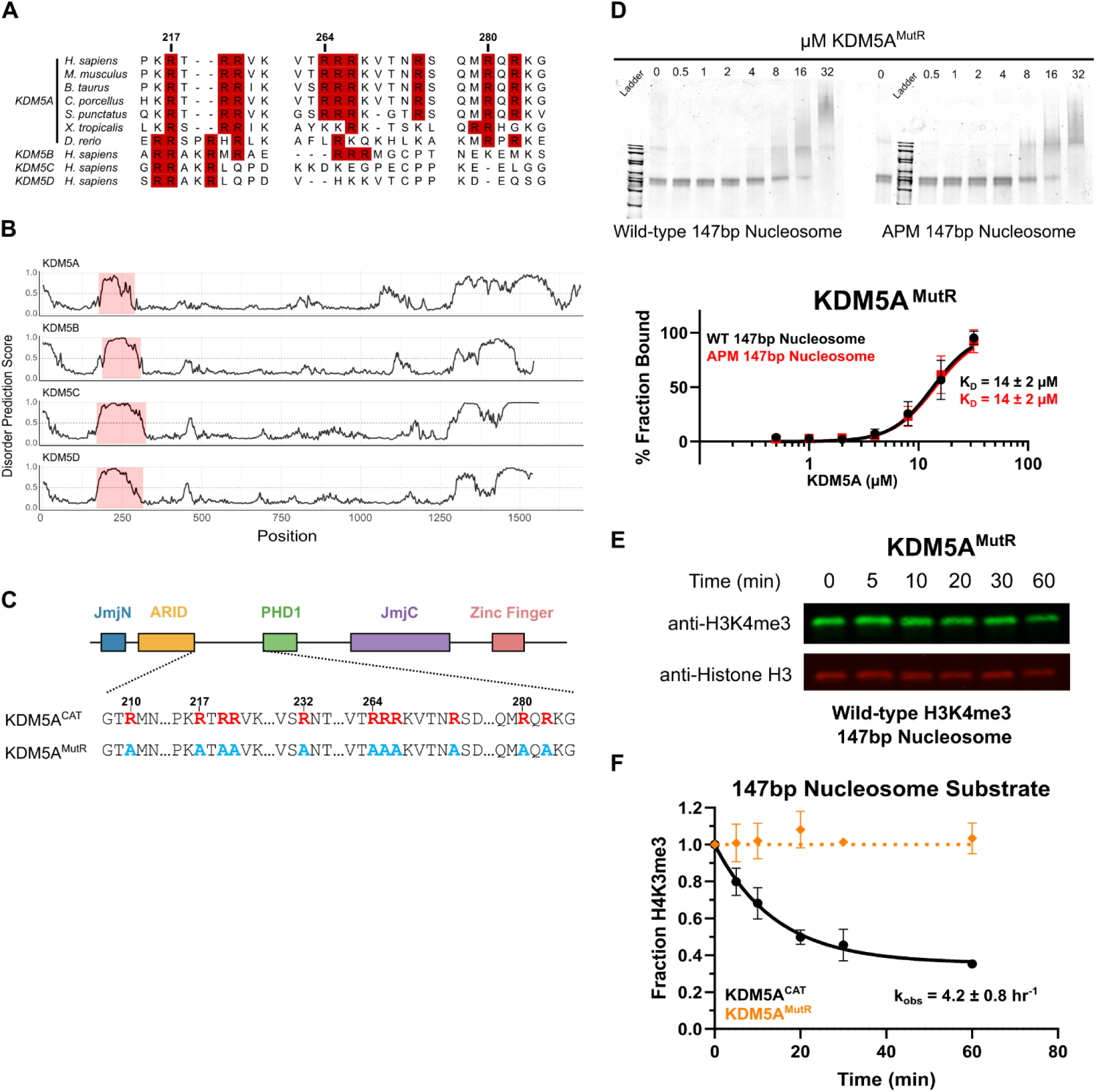
An intrinsically disordered region of KDM5A houses arginine anchors that interact with the H2A/H2B acidic patch. **A.** Multiple sequence alignment of the region between the ARID and PHD1 domains across KDM5A orthologs in vertebrates and KDM5 family proteins in humans. Arginine residues are highlighted in red, alignment performed using ClustalW. **B.** Bioinformatic predictions of disorder in KDM5 family proteins, generated by AIUPred. A score greater than 0.5 indicates disorder. **C.** Schematic of wild-type and arginine mutant KDM5A^MutR^ catalytic constructs. Arginine residues are colored red, and inactivating alanine mutations are colored blue. **D.** EMSA binding assay of arginine inactivating mutant KDM5A^MutR^ to wild-type and acidic patch mutant nucleosomes. Data are presented as the mean ± s.d. from five replicates collected across two independent experiments. **E.** Demethylation of H3K4me3 wild-type 147bp nucleosomes by KDM5A^MutR^, measured by western blot under single turnover conditions (5 µM KDM5A^CAT^, 50 nM nucleosome substrate; n=2). **F.** Demethylation of H3K4me3 wild-type 147bp nucleosomes by KDM5A^CAT^ and KDM5A^MutR^ (From Figure 1D and 2E, respectively).

To assay the function of these arginine residues, we generated a KDM5A construct where all arginine residues between the ARID and PHD1 domains of KDM5A^CAT^ were mutated to alanine (referred to as KDM5A^MutR^) (Figure 2C). Unlike KDM5A^CAT^, which shows 3-fold higher affinity for wild-type vs APM 147bp nucleosomes (Figure 1C), KDM5A^MutR^ displays no preference for wild-type vs APM 147bp nucleosomes (Figure 2D). The similar affinity of KDM5A^MutR^ towards both nucleosomes, which closely matches that of the wild-type enzyme for APM nucleosomes, indicates that the arginine residues between the ARID and PHD1 domains drive acidic patch interactions in KDM5A.

We then assayed the catalytic activity of KDM5A^MutR^ on wild-type 147bp nucleosomes under single turnover conditions and saw no measurable demethylase activity (Figure 2E, 2F). This contrasts with demethylation of 21mer H3K4me3 peptide substrates, where KDM5A^MutR^ demonstrated robust catalytic activity comparable to wild-type KDM5A^CAT^ (Supplementary Figure S5), demonstrating that KDM5A^MutR^ retains its intrinsic catalytic activity.

### Arginine-rich motifs regulate binding to the acidic patch and demethylation catalysis

To identify the specific contributions of the three identified arginine-rich motifs in KDM5A, we expressed and purified KDM5A catalytic constructs where individual arginine motifs are mutated to alanine (Figure 3A), and assayed binding of each to wild-type and APM 147bp nucleosomes. Compared to the WT KDM5A^CAT^, all three arginine motif mutants displayed a reduced affinity to wild-type nucleosomes (Figure 1C, 3B, 3C, Supplementary Figure S6). In contrast, changes to the affinity towards APM nucleosomes were modest, suggesting that all three arginine-rich motifs contribute to acidic patch interaction and affinity.

**Figure 3:**
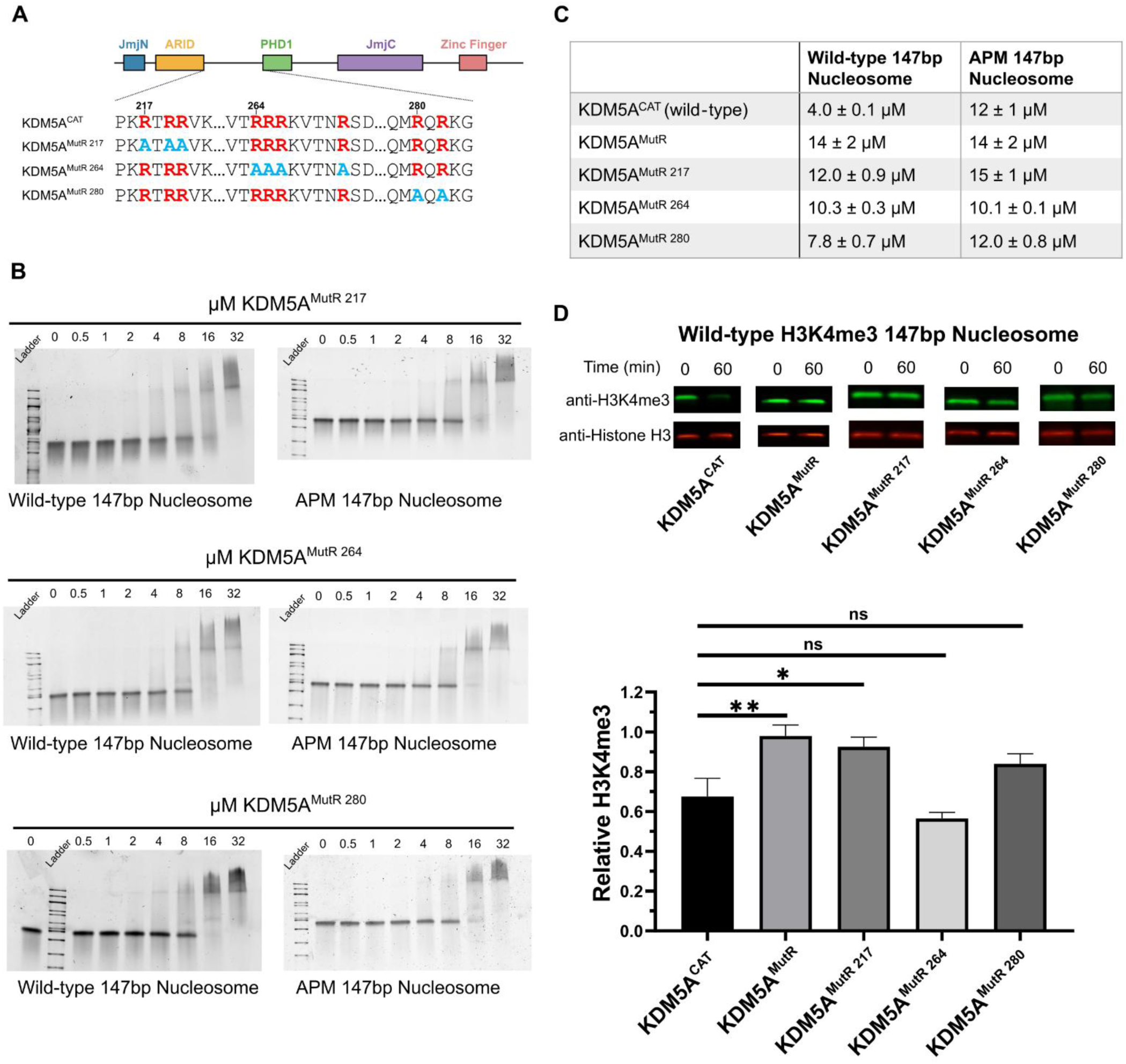
Interactions with the acidic patch and catalytic activity are regulated by multiple arginine-rich motifs in the IDR of KDM5A. **A.** Schematic of wild-type and arginine motif mutant KDM5A catalytic constructs. Arginine residues are colored red, and alanine mutations are colored blue. **B.** EMSA binding assays of KDM5A arginine motif mutants to wild-type and acidic patch mutant nucleosomes. For KDM5A^MutR 217^ and KDM5A^MutR 264^, images are representative of four replicates collected across two independent experiments. For KDM5A^MutR 280^, images are representative of five replicates collected across three independent experiments. Quantification of data is provided in Supplementary Figure S6. **C.** Dissociation constants (K_D_) of KDM5A catalytic constructs bound to wild-type and acidic patch mutant nucleosomes. **D.** Demethylation of H3K4me3 wild-type nucleosomes by KDM5A catalytic constructs, measured by western blot under single turnover conditions (5 µM KDM5A^CAT^, 50 nM nucleosome substrate; ns: p > 0.05; ✱: p ≤ 0.05; ✱✱: p ≤ 0.01; n=3).

To evaluate the impact of the individual arginine-rich motifs on the catalytic activity of KDM5A, we performed demethylation assays on H3K4me3 wild-type 147bp nucleosome with KDM5A^MutR^ as well as all individual patch mutants via western blot at a single time point of 60 minutes. KDM5A^MutR^ and mutations of the arginine motif at residues 217 showed a substantial reduction in catalytic activity (Figure 3D, Supplementary Table S2A). A mutation of the motif at position 280 does suggest a partial loss of catalytic activity, however, the change is not statistically significant according to a student’s t-test (p = 0.052). Interestingly, under assay conditions, mutation of the arginine motif at position 264 showed no measurable loss in catalytic activity despite the deficiency in nucleosome binding (Figure 3C, 3D), indicating that changes in KDM5A catalytic activity are not solely driven by affinity.

### Flanking DNA enhances catalytically productive binding of KDM5A

The shared IDRs of KDM5 family proteins are basic in nature, with KDM5A carrying the highest charge with a pI of 10.2 (Supplementary Table S3). Given that highly basic IDRs have displayed DNA binding capability in their interactions with nucleosomes [45–47], we set out to investigate potential DNA binding activity in the IDR of KDM5A.

We first probed for DNA interaction through the addition of symmetrical 19bp flanking DNA sequences to the nucleosome core particle. Flanking DNA enhanced binding of both KDM5A^CAT^ and KDM5A^MutR^ to wild-type 185bp nucleosome (Figure 4A). We then assayed binding of KDM5A^CAT^ and KDM5A^MutR^ to APM 185bp nucleosome. Unlike with 147bp nucleosome, we observed no loss in affinity on APM 185bp nucleosomes (Supplementary Figure S7), indicating that the addition of flanking DNA can compensate for the loss in affinity upon the elimination of acidic patch binding. To interrogate how improved affinity impacts catalysis, we tested demethylation of 185bp nucleosomes by the KDM5A^CAT^ and KDM5A^MutR^ demethylases under single turnover conditions. While improved affinity to the 185bp nucleosome failed to rescue KDM5A^MutR^ demethylase activity (Figure 4B, Supplementary Table S2B), addition of flanking DNA enhanced the demethylase activity of KDM5A^CAT^ on both wild-type and APM nucleosomes (Figure 1D, 4C).

**Figure 4:**
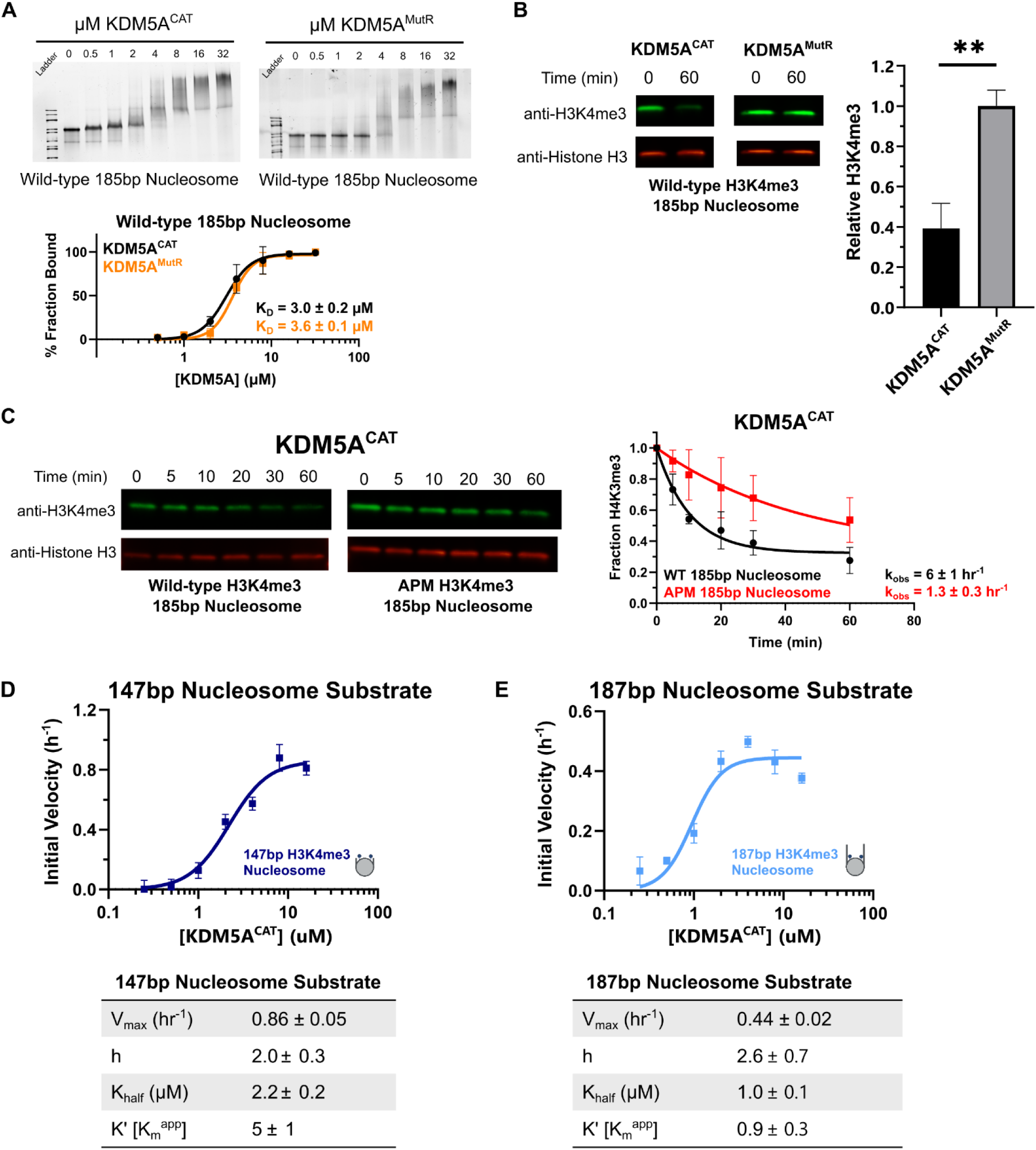
Flanking DNA recognition by KDM5A improves affinity for nucleosome substrate and reduces the K_m_^app^ in catalytic assays. **A.** EMSA binding assay of wild-type KDM5A^CAT^ and KDM5A^MutR^ to wild-type mutant nucleosomes with extranucleosomal flanking DNA. Data are presented as the mean ± s.d. from three replicates collected across two independent experiments. **B.** Demethylation of H3K4me3 nucleosomes with extranucleosomal flanking DNA by wild-type and MutR KDM5A catalytic constructs, measured by western blot. (5 µM KDM5A^CAT^, 50 nM nucleosome substrate; ✱✱: p ≤ 0.01; n=3)**. C.** KDM5A^CAT^ demethylation of H3K4me3 wild-type and APM 185bp nucleosomes, measured by western blot under single turnover conditions (5 µM KDM5A^CAT^, 50 nM nucleosome substrate; n=2). **D, E.** KDM5A^CAT^ demethylation of (**D**) 147bp DNA and (**E**) 187bp DNA H3K4me3 nucleosomes measured by TR-FRET. Initial rates were fitted to an allosteric sigmoidal kinetics model (n=3).

To investigate how flanking DNA improves nucleosome demethylation by KDM5A^CAT^, we monitored demethylation of 147bp and 187bp H3K4me3 nucleosomes using a TR-FRET-based assay. This assay detects the rate of H3K4me2/1 formation under single turnover conditions. Compared to 147bp nucleosomes, demethylation of 187bp nucleosomes shows an approximate 2-fold reduction in V_max_, alongside a more significant 5-fold reduction in the K_m_^app^ (Figure 4D, 4E). Our findings support a model where flanking DNA promotes catalytically productive binding of KDM5A to nucleosome substrate.

### Arginine-rich motifs in the intrinsically disordered region of KDM5A bind to DNA

To evaluate changes in protein conformation and dynamics upon DNA binding and identify DNA binding regions, we performed quantitative cross-linking mass spectrometry. We employed a lysine-directed crosslinker BS3, bis(sulfosuccinimidyl)suberate, to generate intra-protein cross-links on KDM5A, with and without a double-stranded DNA sequence previously identified to interact with the ARID domain, hereon referred to as ARID-C1 (Supplementary Table S4) [6]. Digestion of cross-linking reactions followed by isobaric tag-based quantitative mass spectrometry allows us to observe changes in protein conformation upon DNA binding. Our findings revealed numerous transient intramolecular interactions by the IDR, both within the IDR and with other domains of KDM5A (Figure 5A, Supplementary Figure S8, Supplementary Table S5), consistent with previous observations of proteins engaged in conformationally heterogeneous “fuzzy” interactions [48,49].

**Figure 5:**
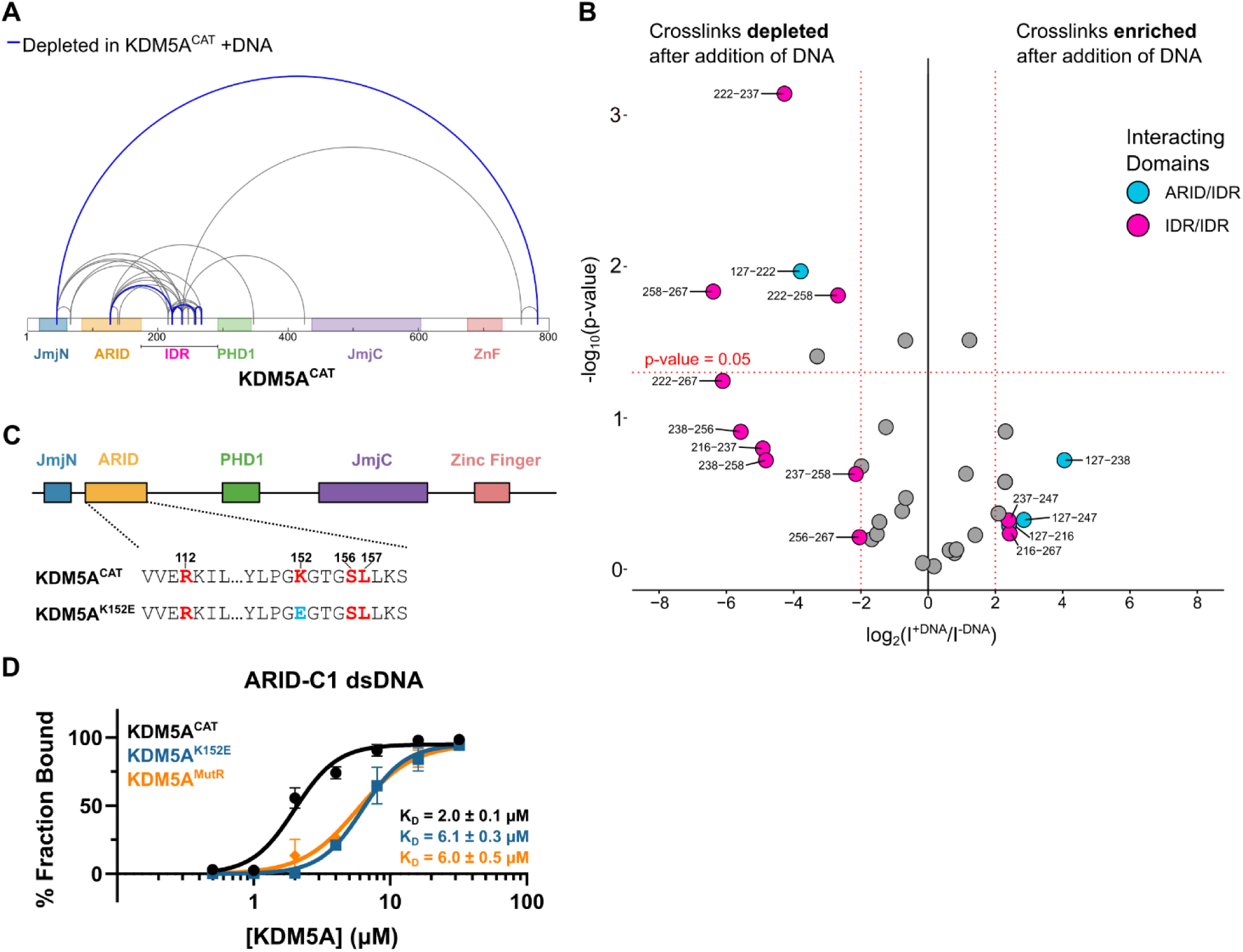
The ARID domain and arginine residues in the intrinsically disordered region of KDM5A bind to free double-stranded DNA *in vitro*. **A.** Map of quantified cross-links in MS analysis of KDM5A^CAT^, ±ARID-C1 dsDNA. Crosslinks with a similar abundance are colored grey, crosslinks depleted by at least four-fold and with a p-value <0.05 are highlighted in blue. **B.** Volcano plot of quantified cross-links, where the log2 Intensity signal ratios of +DNA/-DNA are plotted against the negative log10 of the p-value. The horizontal dashed line marks a p-value of 0.05. Vertical dashed lines mark the four-fold enrichment and depletion thresholds. Cross-links that are beyond the thresholds and are within the ARID and IDR domains are colored: cross-links between the ARID domain and IDR are blue, and cross-links within the IDR are pink. Points are labelled with the positions of the lysine residues cross-linked. **C**. Schematic of wild-type and ARID domain mutant KDM5A^K152E^ catalytic constructs. DNA binding residues are colored red, the inactivating mutation is colored blue. **D.** EMSA binding assay of wild-type KDM5A^CAT^, KDM5A^MutR^, and KDM5A^K152E^ to double-stranded ARID-C1 DNA. For KDM5A^CAT^ and KDM5A^K152E^, data are presented as the mean ± s.d. from three replicates collected across two independent experiments. For KDM5A^MutR^, data are presented as the mean ± s.d. from four replicates collected across two independent experiments.

The analysis of crosslinking data revealed a striking depletion of IDR/IDR interactions upon the addition of DNA (Figure 5B). To further investigate this observation, we measured the affinity of the ARID-C1 DNA for the following constructs: KDM5A^CAT^ (Figure 1A), KDM5A^MutR^ (Figure 2C), and KDM5A^K152E^, a construct where the ARID domain of KDM5A contained a mutation, K152E, previously demonstrated to ablate ARID’s DNA binding ability (Figure 5C) [6]. Compared to the KDM5A^CAT^, both the MutR and K152E mutant proteins showed a similar, approximately three-fold loss in affinity for DNA (Figure 5D, Supplementary Figure S9). These findings suggest that, in addition to binding to the acidic patch on the nucleosome, arginine residues in the IDR of KDM5A bind to DNA, supporting a bifunctional role of the IDR.

### AlphaFold3 simulations predict multivalent interactions of KDM5A with the nucleosome

We next used structural predictions generated by the software AlphaFold3 [50] to test if deep learning forecasts recapitulate the results observed in our biochemical assays. Broadly, the generated predicted structures of KDM5A^CAT^ bound to 187bp nucleosome are consistent with our biochemical observations. All structural predictions featured interactions of arginine anchors in the IDR with the H2A/H2B acidic patch (Figure 6A). Three of five structural predictions generated by AlphaFold3 positioned Arg 280 in the location of the canonical arginine anchor; in the remaining two, Arg 217 was poised as the canonical arginine anchor (Figure 6B, C). Predicted structures also feature core DNA interactions with IDR (Lys 238, Figure 6D) and the ARID domain (Figure 6A, E). We then sought to apply more stringent restrictions upon AlphaFold3 simulations to account for the possibility of false positives. While a prior *in silico* screen developed for acidic patch interactors performed on AlphaFold-Multimer models did not provide any positive hits for KDM5A, an application of the same screening methods to AlphaFold3 models returned all of the aforementioned acidic patch interactions within the KDM5A IDR (Supplementary Table S6) [40].

**Figure 6:**
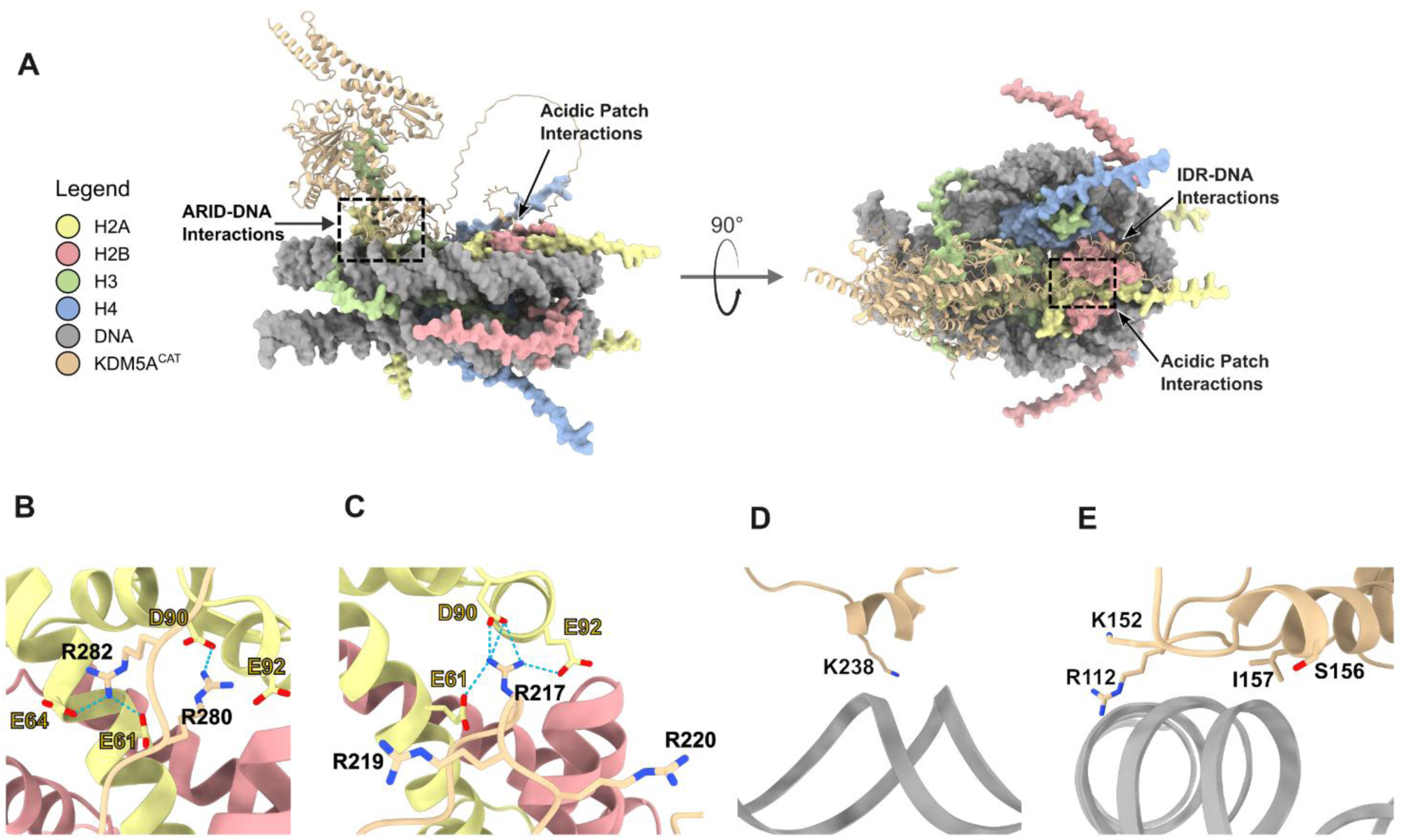
AlphaFold3 Predictions of KDM5A^CAT^ bound to 187bp Nucleosome. **A**. Top-ranked AlphaFold3 prediction of KDM5A^CAT^ interactions with 187bp nucleosome. Interactions of the ARID domain with core DNA and the IDR with the H2A/H2B acidic patch highlighted (Ranking Score = 0.67). **B-E**. Representative AlphaFold3 predictions of (**B-C**) arginine anchors in the IDR interacting with the H2A/H2B acidic patch, (**D**) IDR interactions with nucleosomal core DNA, and (**E**) ARID domain interactions with core DNA. Structures are representative of the variations seen across the five top-ranked models (Ranking Score = 0.67). Atoms of key interacting residues are displayed, annotated, and colored by heteroatom. Potential hydrogen bonding interactions are marked by dashed lines.

## DISCUSSION

Here, we demonstrate that KDM5A demethylation of H3K4me3 is regulated through interactions with the H2A/H2B acidic patch and DNA. In concert with the roles of the PHD1 and PHD3 domains in the regulation of catalysis [3,26,28,51], our findings demonstrate how multivalent interactions with the nucleosome regulate the binding and demethylase activity of KDM5A. A key mediator of these interactions is the intrinsically disordered region between the ARID and PHD1 domains of KDM5A, which interacts with both the H2A/H2B acidic patch and DNA. Our findings also reveal how the plasticity of the IDR of KDM5A enables interactions with multiple features of the nucleosome to regulate catalysis, broadening our understanding of the functional roles of IDRs in chromatin-modifying enzymes.

The presence of acidic patch-interacting arginine residues in the IDR of KDM5A highlights the importance of the ARID-PHD1 cassette separating the two segments of the Jumonji catalytic domain, a feature unique to the KDM5 family. Previous studies have shown the differing roles of the IDR, ARID, and PHD1 domains across KDM5 family proteins [3,26,52,53]. Divergence of these interactions across the KDM5 family likely enables differential regulation of KDM5 family proteins and proper demethylation of their respective targets. While IDR interactions with DNA have yet to be explored in other KDM5s, IDR interactions with both the acidic patch and DNA offer multiple avenues for differential regulation of KDM5 family members on nucleosome substrates.

Loss of acidic patch binding leads to a relatively modest loss in affinity, compared to a more significant loss of activity. This parallels findings observed in other chromatin modifiers, including DOT1L, KDM2A, and SUV420H1, where acidic patch interaction contributed to catalysis not through improved affinity, but through enabling proper positioning of enzymes on the nucleosome substrate [54–58]. The ability of arginine residues within the IDR to also interact with DNA reveals a bifunctional nature previously unobserved in acidic patch-binding chromatin modifiers. Given that inactivating mutations of arginine residues in the IDR of KDM5A lead to a greater loss of demethylation than mutation of the acidic patch in the nucleosome, we propose that arginine-rich motifs in the IDR dynamically interact with both the acidic patch and DNA.

Based on these findings, we hypothesize a model where the IDR promotes catalytically productive binding by enabling proper positioning of KDM5A on the nucleosome through multivalent and conformationally heterogeneous interactions (Figure 7). Arginine-rich motifs in the IDR engage in interactions with the H2A/H2B acidic patch and DNA, adding to a growing list of examples of “fuzzy” interactions between proteins and DNA [46,47,59–61]. Both interactions work in concert to facilitate productive binding to the nucleosome, allowing demethylation to commence. Under this model, the plasticity of these interactions would play a functional role, enabling transient binding to both nucleosomal features. Redundancy of arginine motifs and interaction points on the nucleosome likely safeguards from a loss of demethylation when one interaction is interrupted, consistent with weaker, but still measurable, demethylation activity on APM nucleosomes.

**Figure 7:**
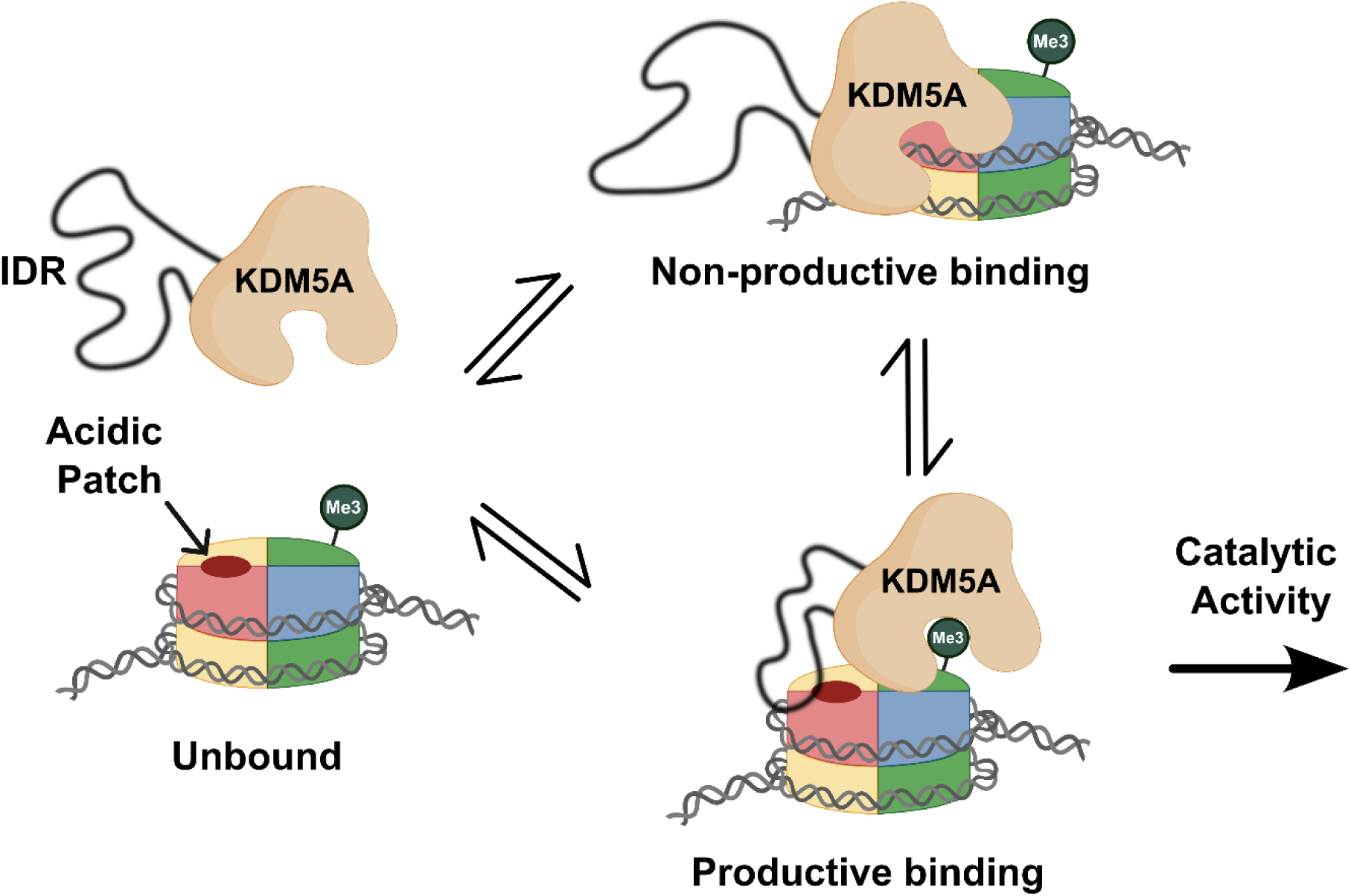
Arginine-rich motifs in the IDR enable catalytically productive binding of KDM5A to nucleosome. KDM5A engages with the nucleosome in heterogeneous conformations. Transient and dynamic interactions of the IDR with the acidic patch and DNA, mediated through arginine-rich motifs in the IDR, are necessary for KDM5A to adopt a catalytically productive binding state. When IDR interactions are disrupted through mutagenesis of arginine-rich motifs, KDM5A binds nucleosomes in a catalytically non-productive manner. (Created in BioRender. Kleinman, J. (2025) https://BioRender.com/o72f887)

Through multiple interaction points on both the enzyme and nucleosome substrates, the IDR of KDM5A offers a dynamic yet specific recognition mechanism for regulating catalytic activity. Our studies highlight how intrinsic disorder in proteins plays a functional role by enabling multivalent modes of binding to work cooperatively. The bifunctional IDR of KDM5A also contributes to the broader regulatory role of the ARID-PHD1 cassette unique to the KDM5 family, a key factor in differentiating the functions of KDM5 family proteins in the context of nucleosome substrate. Our investigation sets the stage for the elucidation of the functions of intrinsically disordered regions of chromatin-modifying enzymes in chromatin recognition and catalysis.

## ACKNOWLEDGEMENTS

We would like to thank Hayden Saunders (University of California, San Francisco) and Ed Linossi, PhD (University of California, San Francisco) for technical assistance.

## AUTHOR CONTRIBUTIONS

Ali Palla: Conceptualization, Formal analysis, Investigation, Methodology, Validation, Visualization, Writing - original draft. Chien-Chu Lin: Investigation, Resources, Writing - review & editing. Michael Trnka: Methodology, Formal analysis, Investigation, Visualization. Emme Leao: Investigation. Nektaria Petronikolou: Resources. Alma Burlingame: Supervision. Robert McGinty: Methodology, Funding acquisition, Supervision, Writing - review & editing. Danica Fujimori: Conceptualization, Funding acquisition, Methodology, Supervision, Writing - review & editing

## DATA AVAILABILITY

Annotated spectra identifying the crosslinks can be viewed at MS-Viewer (https://msviewer.ucsf.edu) with search key: dlhcri59ps

## REFERENCES

[1] R.J. Klose, Q. Yan, Z. Tothova, K. Yamane, H. Erdjument-Bromage, P. Tempst, D.G. Gilliland, Y. Zhang, W.G. Kaelin Jr, The retinoblastoma binding protein RBP2 is an H3K4 demethylase, Cell 128 (2007) 889–900. 10.1016/j.cell.2007.02.013.

[2] J.R. Horton, A. Engstrom, E.L. Zoeller, X. Liu, J.R. Shanks, X. Zhang, M.A. Johns, P.M. Vertino, H. Fu, X. Cheng, Characterization of a linked Jumonji domain of the KDM5/JARID1 family of histone H3 lysine 4 demethylases, J. Biol. Chem. 291 (2016) 2631–2646. 10.1074/jbc.M115.698449.

[3] I.O. Torres, K.M. Kuchenbecker, C.I. Nnadi, R.J. Fletterick, M.J.S. Kelly, D.G. Fujimori, Histone demethylase KDM5A is regulated by its reader domain through a positive-feedback mechanism, Nat. Commun. 6 (2015) 6204. 10.1038/ncomms7204.

[4] A. Kataria, S. Tyagi, Domain architecture and protein-protein interactions regulate KDM5A recruitment to the chromatin, Epigenetics 18 (2023) 2268813. 10.1080/15592294.2023.2268813.

[5] K. Yamane, K. Tateishi, R.J. Klose, J. Fang, L.A. Fabrizio, H. Erdjument-Bromage, J. Taylor-Papadimitriou, P. Tempst, Y. Zhang, PLU-1 is an H3K4 demethylase involved in transcriptional repression and breast cancer cell proliferation, Mol Cell 25 (2007) 801–812. 10.1016/j.molcel.2007.03.001.

[6] S. Tu, Y.-C. Teng, C. Yuan, Y.-T. Wu, M.-Y. Chan, A.-N. Cheng, P.-H. Lin, L.-J. Juan, M.-D. Tsai, The ARID domain of the H3K4 demethylase RBP2 binds to a DNA CCGCCC motif, Nat. Struct. Mol. Biol. 15 (2008) 419–421. 10.1038/nsmb.1400.

[7] E.S. Pilka, T. James, J.H. Lisztwan, Structural definitions of Jumonji family demethylase selectivity, Drug Discov Today 20 (2015) 743–749. 10.1016/j.drudis.2014.12.013.

[8] J. Plch, J. Hrabeta, T. Eckschlager, KDM5 demethylases and their role in cancer cell chemoresistance, Int J Cancer 144 (2019) 221–231. 10.1002/ijc.31881.

[9] G.-J. Yang, M.-H. Zhu, X.-J. Lu, Y.-J. Liu, J.-F. Lu, C.-H. Leung, D.-L. Ma, J. Chen, The emerging role of KDM5A in human cancer, J Hematol Oncol 14 (2021) 30. 10.1186/s13045-021-01041-1.

[10] A. Roesch, B. Becker, S. Meyer, P. Wild, C. Hafner, M. Landthaler, T. Vogt, Retinoblastoma-binding protein 2-homolog 1: a retinoblastoma-binding protein downregulated in malignant melanomas, Mod Pathol 18 (2005) 1249–1257. 10.1038/modpathol.3800413.

[11] B. Dai, H. Huang, F. Guan, G. Zhu, Z. Xiao, B. Mao, H. Su, Z. Hu, Histone demethylase KDM5A inhibits glioma cells migration and invasion by down regulating ZEB1, Biomed Pharmacother 99 (2018) 72–80. 10.1016/j.biopha.2018.01.020.

[12] J. Zeng, Z. Ge, L. Wang, Q. Li, N. Wang, M. Björkholm, J. Jia, D. Xu, The histone demethylase RBP2 Is overexpressed in gastric cancer and its inhibition triggers senescence of cancer cells, Gastroenterology 138 (2010) 981–992. 10.1053/j.gastro.2009.10.004.

[13] H.-J. Choi, H.-S. Joo, H.-Y. Won, K.-W. Min, H.-Y. Kim, T. Son, Y.-H. Oh, J.-Y. Lee, G. Kong, Role of RBP2-Induced ER and IGF1R-ErbB Signaling in Tamoxifen Resistance in Breast Cancer, J Natl Cancer Inst 110 (2018). 10.1093/jnci/djx207.

[14] B. Banelli, E. Carra, F. Barbieri, R. Würth, F. Parodi, A. Pattarozzi, R. Carosio, A. Forlani, G. Allemanni, D. Marubbi, T. Florio, A. Daga, M. Romani, The histone demethylase KDM5A is a key factor for the resistance to temozolomide in glioblastoma, Cell Cycle 14 (2015) 3418–3429. 10.1080/15384101.2015.1090063.

[15] Y.-C. Teng, C.-F. Lee, Y.-S. Li, Y.-R. Chen, P.-W. Hsiao, M.-Y. Chan, F.-M. Lin, H.-D. Huang, Y.-T. Chen, Y.-M. Jeng, C.-H. Hsu, Q. Yan, M.-D. Tsai, L.-J. Juan, Histone demethylase RBP2 promotes lung tumorigenesis and cancer metastasis, Cancer Res 73 (2013) 4711–4721. 10.1158/0008-5472.CAN-12-3165.

[16] T. Feng, Y. Wang, Y. Lang, Y. Zhang, KDM5A promotes proliferation and EMT in ovarian cancer and closely correlates with PTX resistance, Mol Med Rep 16 (2017) 3573–3580. 10.3892/mmr.2017.6960.

[17] H. Wu, L. Xu, X. Hu, KDM5A regulates the growth and gefitinib drug resistance against human lung adenocarcinoma cells, 3 Biotech 12 (2022) 1–8. 10.1007/s13205-021-03018-w.

[18] J.D.E. de Rooij, I.H.I.M. Hollink, S.T.C.J.M. Arentsen-Peters, J.F. van Galen, H. Berna Beverloo, A. Baruchel, J. Trka, D. Reinhardt, E. Sonneveld, M. Zimmermann, T.A. Alonzo, R. Pieters, S. Meshinchi, M.M. van den Heuvel-Eibrink, C.M. Zwaan, NUP98/JARID1A is a novel recurrent abnormality in pediatric acute megakaryoblastic leukemia with a distinct HOX gene expression pattern, Leukemia 27 (2013) 2280–2288. 10.1038/leu.2013.87.

[19] J. Miller, R. Hiltenbrand, T. Lamprecht, A. Seth, S. Abdelhamed, I. Iacobucci, C.G. Mullighan, J.M. Klco, NUP98-KDM5A fusion induces hematopoietic cell proliferation and alters myelo-erythropoietic differentiation, Blood 134 (2019) 3775–3775. 10.1182/blood-2019-130768.

[20] J. Domingo-Reinés, R. Montes, A. Garcia-Moreno, A. Gallardo, J.M. Sanchez-Manas, I. Ellson, M. Lamolda, C. Calabro, J.A. López-Escamez, P. Catalina, P. Carmona-Sáez, P.J. Real, D. Landeira, V. Ramos-Mejia, The pediatric leukemia oncoprotein NUP98-KDM5A induces genomic instability that may facilitate malignant transformation, Cell Death Dis 14 (2023) 357. 10.1038/s41419-023-05870-5.

[21] A. Patsialou, D. Wilsker, E. Moran, DNA-binding properties of ARID family proteins, Nucleic Acids Res. 33 (2005) 66–80. 10.1093/nar/gki145.

[22] G.-J. Yang, J. Wu, L. Miao, M.-H. Zhu, Q.-J. Zhou, X.-J. Lu, J.-F. Lu, C.-H. Leung, D.-L. Ma, J. Chen, Pharmacological inhibition of KDM5A for cancer treatment, Eur J Med Chem 226 (2021) 113855. 10.1016/j.ejmech.2021.113855.

[23] N. Petronikolou, J.E. Longbotham, D.G. Fujimori, Extended recognition of the histone H3 tail by histone demethylase KDM5A, Biochemistry 59 (2020) 647–651. 10.1021/acs.biochem.9b01036.

[24] M. Hoekstra, K.K. Biggar, Identification of in vitro JMJD lysine demethylase candidate substrates via systematic determination of substrate preference, Anal Biochem 633 (2021) 114429. 10.1016/j.ab.2021.114429.

[25] M. Hoekstra, N.H. Ridgeway, K.K. Biggar, Characterization of KDM5 lysine demethylase family substrate preference and identification of novel substrates, J Biochem 173 (2022) 31–42. 10.1093/jb/mvac081.

[26] J.E. Longbotham, C.M. Chio, V. Dharmarajan, M.J. Trnka, I.O. Torres, D. Goswami, K. Ruiz, A.L. Burlingame, P.R. Griffin, D.G. Fujimori, Histone H3 binding to the PHD1 domain of histone demethylase KDM5A enables active site remodeling, Nat. Commun. 10 (2019) 94. 10.1038/s41467-018-07829-z.

[27] J.E. Longbotham, M.J.S. Kelly, D.G. Fujimori, Recognition of histone H3 methylation states by the PHD1 domain of histone demethylase KDM5A, ACS Chem. Biol. 18 (2023) 1915–1925. 10.1021/acschembio.0c00976.

[28] G.G. Wang, J. Song, Z. Wang, H.L. Dormann, F. Casadio, H. Li, J.-L. Luo, D.J. Patel, C.D. Allis, Haematopoietic malignancies caused by dysregulation of a chromatin-binding PHD finger, Nature 459 (2009) 847–851. 10.1038/nature08036.

[29] M.Y. Zhang, H. Yang, G. Ortiz, M.J. Trnka, N. Petronikolou, A.L. Burlingame, W.F. DeGrado, D.G. Fujimori, Covalent labeling of a chromatin reader domain using proximity-reactive cyclic peptides, Chem Sci 13 (2022) 6599–6609. 10.1039/d2sc00555g.

[30] R.K. McGinty, S. Tan, Recognition of the nucleosome by chromatin factors and enzymes, Curr. Opin. Struct. Biol. 37 (2016) 54–61. 10.1016/j.sbi.2015.11.014.

[31] R.K. McGinty, S. Tan, Principles of nucleosome recognition by chromatin factors and enzymes, Curr. Opin. Struct. Biol. 71 (2021) 16–26. 10.1016/j.sbi.2021.05.006.

[32] W. Xu, H. Zhang, W. Guo, L. Jiang, Y. Zhao, Y. Peng, Deciphering principles of nucleosome interactions and impact of cancer-associated mutations from comprehensive interaction network analysis, Brief. Bioinform. 25 (2024). 10.1093/bib/bbad532.

[33] N. Petronikolou, J.E. Longbotham, D.G. Fujimori, Dissecting contributions of catalytic and reader domains in regulation of histone demethylation, Methods Enzymol. 639 (2020) 217–236. 10.1016/bs.mie.2020.04.015.

[34] M. Wu, D. Hayward, J.H. Kalin, Y. Song, J.W.R. Schwabe, P.A. Cole, Lysine-14 acetylation of histone H3 in chromatin confers resistance to the deacetylase and demethylase activities of an epigenetic silencing complex, (2018). 10.7554/eLife.37231.

[35] P.N. Dyer, R.S. Edayathumangalam, C.L. White, Y. Bao, S. Chakravarthy, U.M. Muthurajan, K. Luger, Reconstitution of nucleosome core particles from recombinant histones and DNA, Methods Enzymol. 375 (2004) 23–44. 10.1016/s0076-6879(03)75002-2.

[36] A. Skrajna, D. Goldfarb, K.M. Kedziora, E.M. Cousins, G.D. Grant, C.J. Spangler, E.H. Barbour, X. Yan, N.A. Hathaway, N.G. Brown, J.G. Cook, M.B. Major, R.K. McGinty, Comprehensive nucleosome interactome screen establishes fundamental principles of nucleosome binding, Nucleic Acids Res. 48 (2020) 9415–9432. 10.1093/nar/gkaa544.

[37] K. Luger, T.J. Rechsteiner, T.J. Richmond, Preparation of nucleosome core particle from recombinant histones, Methods Enzymol 304 (1999) 3–19. 10.1016/s0076-6879(99)04003-3.

[38] R. Rad, J. Li, J. Mintseris, J. O’Connell, S.P. Gygi, D.K. Schweppe, Improved Monoisotopic Mass Estimation for Deeper Proteome Coverage, Journal of Proteome Research (2020). 10.1021/acs.jproteome.0c00563.

[39] J. Abramson, J. Adler, J. Dunger, R. Evans, T. Green, A. Pritzel, O. Ronneberger, L. Willmore, A.J. Ballard, J. Bambrick, S.W. Bodenstein, D.A. Evans, C.-C. Hung, M. O’Neill, D. Reiman, K. Tunyasuvunakool, Z. Wu, A. Žemgulytė, E. Arvaniti, C. Beattie, O. Bertolli, A. Bridgland, A. Cherepanov, M. Congreve, A.I. Cowen-Rivers, A. Cowie, M. Figurnov, F.B. Fuchs, H. Gladman, R. Jain, Y.A. Khan, C.M.R. Low, K. Perlin, A. Potapenko, P. Savy, S. Singh, A. Stecula, A. Thillaisundaram, C. Tong, S. Yakneen, E.D. Zhong, M. Zielinski, A. Žídek, V. Bapst, P. Kohli, M. Jaderberg, D. Hassabis, J.M. Jumper, Accurate structure prediction of biomolecular interactions with AlphaFold 3, Nature 630 (2024) 493–500. 10.1038/s41586-024-07487-w.

[40] A.M. James, E.W. Schmid, J.C. Walter, L. Farnung, In silico screening identifies SHPRH as a novel nucleosome acidic patch interactor, bioRxiv (2024) 2024.06.26.600687. 10.1101/2024.06.26.600687.

[41] A.J. Barbera, J.V. Chodaparambil, B. Kelley-Clarke, V. Joukov, J.C. Walter, K. Luger, K.M. Kaye, The nucleosomal surface as a docking station for Kaposi’s sarcoma herpesvirus LANA, Science 311 (2006) 856–861. 10.1126/science.1120541.

[42] M. Vinogradova, V.S. Gehling, A. Gustafson, S. Arora, C.A. Tindell, C. Wilson, K.E. Williamson, G.D. Guler, P. Gangurde, W. Manieri, J. Busby, E.M. Flynn, F. Lan, H.-J. Kim, S. Odate, A.G. Cochran, Y. Liu, M. Wongchenko, Y. Yang, T.K. Cheung, T.M. Maile, T. Lau, M. Costa, G.V. Hegde, E. Jackson, R. Pitti, D. Arnott, C. Bailey, S. Bellon, R.T. Cummings, B.K. Albrecht, J.-C. Harmange, J.R. Kiefer, P. Trojer, M. Classon, An inhibitor of KDM5 demethylases reduces survival of drug-tolerant cancer cells, Nat. Chem. Biol. 12 (2016) 531–538. 10.1038/nchembio.2085.

[43] V. Bavetsias, R.M. Lanigan, G.F. Ruda, B. Atrash, M.G. McLaughlin, A. Tumber, N.Y. Mok, Y.-V. Le Bihan, S. Dempster, K.J. Boxall, F. Jeganathan, S.B. Hatch, P. Savitsky, S. Velupillai, T. Krojer, K.S. England, J. Sejberg, C. Thai, A. Donovan, A. Pal, G. Scozzafava, J.M. Bennett, A. Kawamura, C. Johansson, A. Szykowska, C. Gileadi, N.A. Burgess-Brown, F. von Delft, U. Oppermann, Z. Walters, J. Shipley, F.I. Raynaud, S.M. Westaway, R.K. Prinjha, O. Fedorov, R. Burke, C.J. Schofield, I.M. Westwood, C. Bountra, S. Müller, R.L.M. van Montfort, P.E. Brennan, J. Blagg, 8-substituted pyrido[3,4-d]pyrimidin-4(3H)-one derivatives as potent, cell permeable, KDM4 (JMJD2) and KDM5 (JARID1) histone lysine demethylase inhibitors, J. Med. Chem. 59 (2016) 1388–1409. 10.1021/acs.jmedchem.5b01635.

[44] J.R. Horton, C.B. Woodcock, Q. Chen, X. Liu, X. Zhang, J. Shanks, G. Rai, B.T. Mott, D.J. Jansen, S.C. Kales, M.J. Henderson, M. Cyr, K. Pohida, X. Hu, P. Shah, X. Xu, A. Jadhav, D.J. Maloney, M.D. Hall, A. Simeonov, H. Fu, P.M. Vertino, X. Cheng, Structure-based engineering of irreversible inhibitors against histone lysine demethylase KDM5A, J. Med. Chem. 61 (2018) 10588–10601. 10.1021/acs.jmedchem.8b01219.

[45] W.H. Liu, J. Zheng, J.L. Feldman, M.A. Klein, V.I. Kuznetsov, C.L. Peterson, P.R. Griffin, J.M. Denu, Multivalent interactions drive nucleosome binding and efficient chromatin deacetylation by SIRT6, Nat. Commun. 11 (2020) 5244. 10.1038/s41467-020-19018-y.

[46] A.L. Turner, M. Watson, O.G. Wilkins, L. Cato, A. Travers, J.O. Thomas, K. Stott, Highly disordered histone H1-DNA model complexes and their condensates, Proc. Natl. Acad. Sci. U. S. A. 115 (2018) 11964–11969. 10.1073/pnas.1805943115.

[47] M.A. Desai, H.D. Webb, L.M. Sinanan, J.N. Scarsdale, N.M. Walavalkar, G.D. Ginder, D.C. Williams Jr, An intrinsically disordered region of methyl-CpG binding domain protein 2 (MBD2) recruits the histone deacetylase core of the NuRD complex, Nucleic Acids Res. 43 (2015) 3100–3113. 10.1093/nar/gkv168.

[48] J.P. Singh, Y. Li, Y.-Y. Chen, S.-T.D. Hsu, R. Page, W. Peti, T.-C. Meng, The catalytic activity of TCPTP is auto-regulated by its intrinsically disordered tail and activated by Integrin alpha-1, Nat Commun 13 (2022) 94. 10.1038/s41467-021-27633-6.

[49] M. Fuxreiter, Context-dependent, fuzzy protein interactions: Towards sequence-based insights, Curr Opin Struct Biol 87 (2024) 102834. 10.1016/j.sbi.2024.102834.

[50] J. Abramson, J. Adler, J. Dunger, R. Evans, T. Green, A. Pritzel, O. Ronneberger, L. Willmore, A.J. Ballard, J. Bambrick, S.W. Bodenstein, D.A. Evans, C.-C. Hung, M. O’Neill, D. Reiman, K. Tunyasuvunakool, Z. Wu, A. Žemgulytė, E. Arvaniti, C. Beattie, O. Bertolli, A. Bridgland, A. Cherepanov, M. Congreve, A.I. Cowen-Rivers, A. Cowie, M. Figurnov, F.B. Fuchs, H. Gladman, R. Jain, Y.A. Khan, C.M.R. Low, K. Perlin, A. Potapenko, P. Savy, S. Singh, A. Stecula, A. Thillaisundaram, C. Tong, S. Yakneen, E.D. Zhong, M. Zielinski, A. Žídek, V. Bapst, P. Kohli, M. Jaderberg, D. Hassabis, J.M. Jumper, Accurate structure prediction of biomolecular interactions with AlphaFold 3, Nature 630 (2024) 493–500. 10.1038/s41586-024-07487-w.

[51] B.J. Klein, L. Piao, Y. Xi, H. Rincon-Arano, S.B. Rothbart, D. Peng, H. Wen, C. Larson, X. Zhang, X. Zheng, M.A. Cortazar, P.V. Peña, A. Mangan, D.L. Bentley, B.D. Strahl, M. Groudine, W. Li, X. Shi, T.G. Kutateladze, The histone-H3K4-specific demethylase KDM5B binds to its substrate and product through distinct PHD fingers, Cell Rep 6 (2014) 325–335. 10.1016/j.celrep.2013.12.021.

[52] Y. Zhang, H. Yang, X. Guo, N. Rong, Y. Song, Y. Xu, W. Lan, X. Zhang, M. Liu, Y. Xu, C. Cao, The PHD1 finger of KDM5B recognizes unmodified H3K4 during the demethylation of histone H3K4me2/3 by KDM5B, Protein Cell 5 (2014) 837–850. 10.1007/s13238-014-0078-4.

[53] F.S. Ugur, M.J.S. Kelly, D.G. Fujimori, Chromatin sensing by the auxiliary domains of KDM5C regulates its demethylase activity and is disrupted by X-linked intellectual disability mutations, J. Mol. Biol. 435 (2023) 167913. 10.1016/j.jmb.2022.167913.

[54] M.I. Valencia-Sánchez, P. De Ioannes, M. Wang, N. Vasilyev, R. Chen, E. Nudler, J.-P. Armache, K.-J. Armache, Structural Basis of Dot1L Stimulation by Histone H2B Lysine 120 Ubiquitination, Mol Cell 74 (2019) 1010–1019.e6. 10.1016/j.molcel.2019.03.029.

[55] C.J. Anderson, M.R. Baird, A. Hsu, E.H. Barbour, Y. Koyama, M.J. Borgnia, R.K. McGinty, Structural Basis for Recognition of Ubiquitylated Nucleosome by Dot1L Methyltransferase, Cell Rep 26 (2019) 1681–1690.e5. 10.1016/j.celrep.2019.01.058.

[56] C.J. Spangler, A. Skrajna, C.A. Foley, A. Nguyen, G.R. Budziszewski, D.N. Azzam, E.C. Arteaga, H.C. Simmons, C.B. Smith, N.A. Wesley, E.M. Wilkerson, J.-M.E. McPherson, D. Kireev, L.I. James, S.V. Frye, D. Goldfarb, R.K. McGinty, Structural basis of paralog-specific KDM2A/B nucleosome recognition, Nat Chem Biol 19 (2023) 624–632. 10.1038/s41589-023-01256-y.

[57] L. Huang, Y. Wang, H. Long, H. Zhu, Z. Wen, L. Zhang, W. Zhang, Z. Guo, L. Wang, F. Tang, J. Hu, K. Bao, P. Zhu, G. Li, Z. Zhou, Structural insight into H4K20 methylation on H2A.Z-nucleosome by SUV420H1, Mol Cell 83 (2023) 2884–2895.e7. 10.1016/j.molcel.2023.07.001.

[58] C.J. Spangler, S.P. Yadav, D. Li, C.N. Geil, C.B. Smith, G.G. Wang, T.-H. Lee, R.K. McGinty, DOT1L activity in leukemia cells requires interaction with ubiquitylated H2B that promotes productive nucleosome binding, Cell Rep 38 (2022) 110369. 10.1016/j.celrep.2022.110369.

[59] R. Pricer, J.E. Gestwicki, A.K. Mapp, From Fuzzy to Function: The New Frontier of Protein-Protein Interactions, Acc Chem Res 50 (2017) 584–589. 10.1021/acs.accounts.6b00565.

[60] E. Komives, R. Sanchez-Rodriguez, H. Taghavi, M. Fuxreiter, Fuzzy protein-DNA interactions and beyond: A common theme in transcription?, Curr Opin Struct Biol 89 (2024) 102941. 10.1016/j.sbi.2024.102941.

[61] C. Langini, A. Caflisch, A. Vitalis, The ATAD2 bromodomain binds different acetylation marks on the histone H4 in similar fuzzy complexes, The Journal of Biological Chemistry 292 (2017). 10.1074/jbc.M117.786350.

