## Supplementary Tables S1-S6 for "An intrinsically disordered region of histone demethylase KDM5A activates catalysis through interactions with the nucleosomal acidic patch and DNA"

### Supplementary Table S1

Protein sequences used in generation of AlphaFold3 structures of KDM5A$^{CAT}$ bound to 187bp nucleosome.

| Type | Copies | Identifier | Sequence |
|---|---|---|---|
| Protein | 2 | Histone H2A | SGRGKQGGKARAKAKTRSSRAGLQFPVGRVHRLLRKGNYAE<br>RVGAGAPVYLAAVLEYLTAEILELAGNAARDNKKTRIIPRHLQL<br>AIRNDEELNKLLGKVTIAQGGVLPNIQAVLLPKKTESHHKAKGK |
| Protein | 2 | Histone H2B | PEPAKSAPAPKKGSKKAVTKAQKKDGKKRKRSRKESYSVYVY<br>KVLKQVHPDTGISSKAMGIMNSFVNDIFERIAGEASRLAHYNK<br>RSTITSREIQTAVRLLLPGELAKHAVSEGTKAVTKYTSSK |
| Protein | 2 | Histone H3.2 | ARTKQTARKSTGGKAPRKQLATKAARKSAPATGGVKKPHRYR<br>PGTVALREIRRYQKSTELLIRKLPFQRLVREIAQDFKTDLRFQS<br>SAVMALQEASEAYLVGLFEDTNLCAIHAKRVTIMPKDIQLARRI<br>RGERA |
| Protein | 2 | Histone H4 | SGRGKGGKGLGKGGAKRHRKVLRDNIQGITKPAIRRLARRGG<br>VKRISGLIYEETRGVLKVFLENVIRDAVTYTEHAKRKTVTAMDV<br>VYALKRQGRTLYGFGG |
| Protein | 1 | KDM5A$^{CAT}$ (1-801) | MAGVGPGGYAAEFVPPPECPVFEPSWEEFTDPLSFIGRIRPLA<br>EKTGICKIRPPKDWQPPFACEVKSFRFTPRVQRLNELEAMTRV<br>RLDFLDQLAKFWELQGSTLKIPVVERKILDLYALSKIVASKGGF<br>EMVTKEKKWSKVGSRLGYLPGKGTGSLLKSHYERILYPYELF<br>QSGVSLMGVQMPNLDLKEKVEPEVLSTDTQTSPEPGTRMNIL<br>PKRTRRVKTQSESGDVSRNTELKKLQIFGAGPKVVGLAMGTK<br>DKEDEVTRRRKVTNRSDAFNMQMRQRKGTLSVNFVDLYVCM<br>FCGRGNNEDKLLLCDGCDDSYHTFCLIPPLPDVPKGDWRCPK<br>CVAEECSKPREAFGFEQAVREYTLQSFGEMADNFKSDYFNM<br>PVHMVPTELVEKEFWRLVSSIEEDVIVEYGADISSKDFGSGFP<br>VKDGRRKILPEEEEYALSGWNLNNMPVLEQSVLAHINVDISGM<br>KVPWLYVGMCFSSFCWHIEDHWSYSINYLHWGEPKTWYGVP<br>SHAAEQLEEVMRELAPELFESQPDLLHQLVTIMNPNVLMEHG<br>VPVYRTNQCAGEFVVTFPRAYHSGFNQGYNFAEAVNFCTAD<br>WLPIGRQCVNHYRRLRRHCVFSHEELIFKMAADPECLDVGLA<br>AMVCKELTLMTEEETRLRESVVQMGVLMSEEEVFELVPDDER<br>QCSACRTTCFLSALTCSCNPERLVCLYHPTDLCPCPMQKKCL<br>RYRYPLEDLPSLLYGVKVRAQSYDTWVSRVTEALSANFNHKK<br>DLIELRVMLEDAEDRKYPENDLFRKLRDAVKEAETCASVAQLL |
| DNA | 1 | 187bp DNA (+strand) | GCGGTGGCGGCCGCTCTAGAACAGGATGTATATATCTGACA<br>CGTGCCTGGAGACTAGGGAGTAATCCCCTTGGCGGTTAAAA<br>CGCGGGGGACAGCGCGTACGTGCGTTTAAGCGGTGCTAGA<br>GCTGTCTACGACCAATTGAGCGGCCTCGGCACCGGGATTC<br>TCCAGGGCGGCCGCGTATAGGGTCC |
| DNA | 1 | 187bp DNA (-strand) | GGACCCTATACGCGGCCGCCCTGGAGAATCCCGGTGCCGA<br>GGCCGCTCAATTGGTCGTAGACAGCTCTAGCACCGCTTAAA<br>CGCACGTACGCGCTGTCCCCCGCGTTTTAACCGCCAAGGG<br>GATTACTCCCTAGTCTCCAGGCACGTGTCAGATATATACATC<br>CTGTTCTAGAGCGGCCGCCACCGC |

### Supplementary Table S2

**A**

| | **Relative H3K4me3** |
|---|---|
| KDM5A$^{CAT}$ | 0.68 ± 0.09 |
| KDM5A$^{MutR}$ | 0.98 ± 0.06 |
| KDM5A$^{MutR\ 217}$ | 0.92 ± 0.05 |
| KDM5A$^{MutR\ 264}$ | 0.56 ± 0.03 |
| KDM5A$^{MutR\ 280}$ | 0.84 ± 0.05 |

**B**

| | **Relative H3K4me3** |
|---|---|
| KDM5A$^{CAT}$ | 0.4 ± 0.1 |
| KDM5A$^{MutR}$ | 1.00 ± 0.08 |

**A.** Demethylation of H3K4me3 wild-type 147bp nucleosomes by KDM5A catalytic constructs, measured by western blot, as shown in Figure 3D
**B.** Demethylation of H3K4me3 nucleosomes with extranucleosomal flanking DNA (185bp Nucleosome) by KDM5A$^{CAT}$ and KDM5A$^{MutR}$, measured by western blot, as shown in Figure 4B

### Supplementary Table S3

pI values for the disordered region between the ARID and PHD1 domains of KDM5 family proteins in humans.

| | Start | End | pI |
|---|---|---|---|
| KDM5A | 177 | 289 | 10.21 |
| KDM5B | 188 | 308 | 9.34 |
| KDM5C | 170 | 325 | 8.69 |
| KDM5D | 170 | 325 | 9.22 |

**Supplementary Table S4**

Sequence of ARID domain binding ARID-C1 double-stranded DNA (Tu et al. 2008). Previously characterized ARID domain recognition sequence is underlined

**ARID-C1 double-stranded DNA**

| | |
|---|---|
| ARID-C1 Fwd | GGGCTCCCGCCCCACGAAAAG |
| ARID-C1 Rev | CTTTTCGTGGGGCGGGAGCCC |

### Supplementary Table S5

| Xlink.AA.1 | Xlink.AA.2 | mean.Intensity | avg.foldChange | std.dev | p.value |
|---|---|---|---|---|---|
| 45 | 216 | 49.70668 | 1.123849 | 1.153159 | 0.233462 |
| 45 | 222 | 50.47023 | -0.66074 | 0.914488 | 0.337304 |
| 45 | 238 | 49.3832 | -0.16064 | 2.086111 | 0.906109 |
| 45 | 66 | 48.27565 | 0.841318 | 3.787169 | 0.737467 |
| 45 | 783 | 51.13895 | -3.29929 | 1.16498 | 0.039136 |
| 66 | 222 | 50.90028 | 2.093052 | 3.659465 | 0.426263 |
| 66 | 238 | 49.98761 | 2.289906 | 2.578368 | 0.263831 |
| 127 | 138 | 51.89631 | -0.67796 | 0.210897 | 0.030775 |
| 127 | 216 | 54.08379 | 2.406626 | 5.285249 | 0.512937 |
| 127 | 222 | 49.18562 | -3.79081 | 0.686199 | 0.010747 |
| 127 | 238 | 54.59479 | 4.053702 | 3.590077 | 0.189664 |
| 127 | 247 | 51.143 | 2.848083 | 5.585552 | 0.470306 |
| 127 | 267 | 49.06106 | -1.68048 | 5.189744 | 0.63135 |
| 127 | 347 | 50.3744 | 0.174865 | 4.509431 | 0.952561 |
| 141 | 238 | 51.79916 | 0.785082 | 4.258213 | 0.779741 |
| 216 | 222 | 54.20331 | -1.25663 | 0.809152 | 0.114874 |
| 216 | 237 | 43.65031 | -4.91627 | 3.872157 | 0.15891 |
| 216 | 238 | 53.99073 | 0.637121 | 2.964647 | 0.745464 |
| 216 | 258 | 49.52871 | 1.401406 | 3.843597 | 0.592256 |
| 216 | 267 | 50.00598 | 2.42715 | 6.385353 | 0.577953 |
| 222 | 237 | 47.9504 | -4.27359 | 0.199017 | 0.000722 |
| 222 | 238 | 53.8817 | -1.52271 | 4.080279 | 0.584303 |
| 222 | 256 | 46.88212 | -1.44937 | 2.954224 | 0.484955 |
| 222 | 258 | 48.01261 | -2.68088 | 0.585092 | 0.015509 |
| 222 | 267 | 44.00502 | -6.10936 | 2.63709 | 0.056861 |
| 237 | 247 | 58.82271 | 2.390749 | 4.715925 | 0.472516 |
| 237 | 258 | 42.73482 | -2.1476 | 2.208586 | 0.234176 |
| 238 | 256 | 51.82255 | -5.57302 | 3.743277 | 0.123201 |
| 238 | 258 | 47.46376 | -4.82472 | 4.277194 | 0.189942 |
| 238 | 425 | 46.09023 | -1.98198 | 1.872927 | 0.208272 |
| 238 | 758 | 46.1863 | 2.302102 | 1.541741 | 0.122607 |
| 247 | 258 | 55.33451 | -0.77243 | 1.299594 | 0.411472 |
| 256 | 267 | 49.75326 | -2.04077 | 5.927471 | 0.611461 |
| 258 | 267 | 45.29991 | -6.39175 | 1.353181 | 0.014613 |
| 758 | 783 | 50.47841 | 1.228887 | 0.381314 | 0.030627 |

Cross-linked residues identified using BS3 cross-linking coupled with mass spectrometry, as displayed in Figure 5A, 5B and Supplementary Figure S8. The cross-link (XLink.AA) values indicate the amino acid number of the KDM5A protein sequence. Average intensities, average fold changes, standard deviations, and p-values are calculated from experiments performed in triplicate

### Supplementary Table S6

Acidic patch interacting residues by *in silico* screen developed by James et al. on structures of KDM5A$^{CAT}$ on a 187bp nucleosome generated by AlphaFold3

| Model Number | Model Confidence Score | Residue | Residue Index | Sequence Context | Distance (Å) | Residue pLDDT |
|---|---|---|---|---|---|---|
| 0 | 0.67 | Arg | 280 | NRSDAFNMQM_QRKGTLSVNF | 0.64 | 57.15 |
| 0 | 0.67 | Arg | 282 | SDAFNMQMRQ_KGTLSVNFVD | 2.06 | 49.26 |
| 1 | 0.67 | Arg | 280 | NRSDAFNMQM_QRKGTLSVNF | 0.75 | 55.96 |
| 1 | 0.67 | Arg | 282 | SDAFNMQMRQ_KGTLSVNFVD | 5.79 | 48.38 |
| 2 | 0.67 | Arg | 280 | NRSDAFNMQM_QRKGTLSVNF | 0.5 | 59.85 |
| 2 | 0.67 | Arg | 282 | SDAFNMQMRQ_KGTLSVNFVD | 4.44 | 51.28 |
| 3 | 0.67 | Arg | 217 | PGTRMNILPK_TRRVKTQSES | 0.99 | 37.88 |
| 4 | 0.67 | Arg | 217 | PGTRMNILPK_TRRVKTQSES | 0.66 | 39.07 |
