## Supplementary Figures S1-S9 for "An intrinsically disordered region of histone demethylase KDM5A activates catalysis through interactions with the nucleosomal acidic patch and DNA"

### Supplementary Figure S1

**A**

$LANA_{1\text{-}23}$ MAPPGMRL**R**SGRSTGAPLTRGSC

$LANA_{1\text{-}23}$ R9A MAPPGMRL**A**SGRSTGAPLTRGSC

**B**

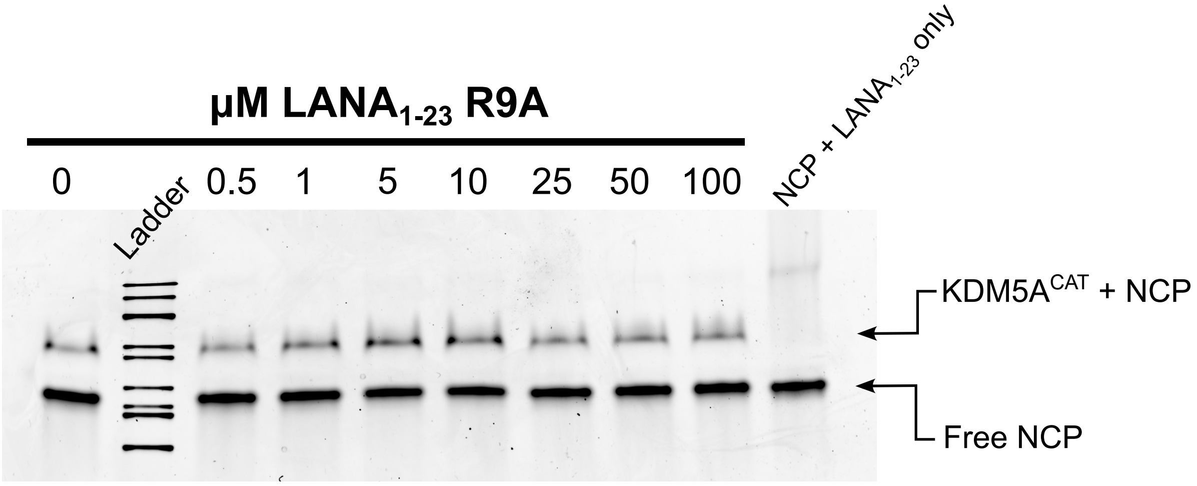


**A.** Schematic of $LANA_{1\text{-}23}$ wild-type and R9A mutant peptides used in competition binding assays

**B.** EMSA competition assay between $KDM5A^{CAT}$, $LANA_{1\text{-}23}$ R9A peptide, and nucleosome. Image is representative of assay performed in triplicate. Binding reactions were prepared with 8 µM KDM5A and 100 nM wild-type nucleosome, then competed with increasing amounts of LANA peptide

### Supplementary Figure S2

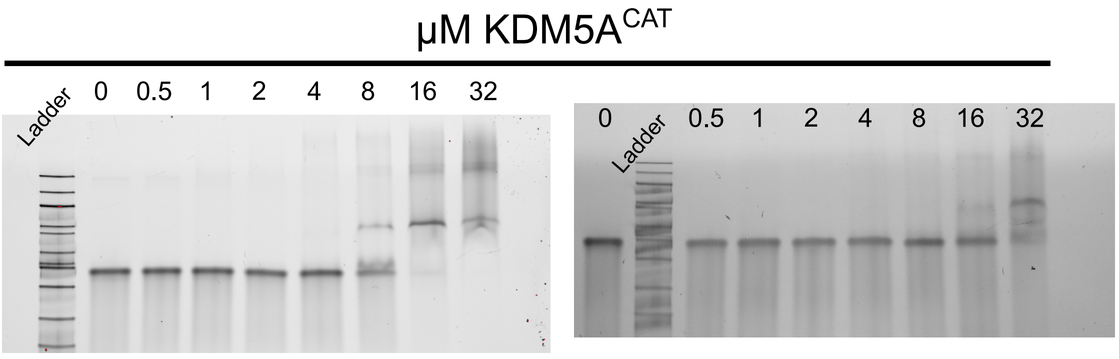


Wild-type H3K4me3
147bp Nucleosome

APM H3K4me3
147bp Nucleosome

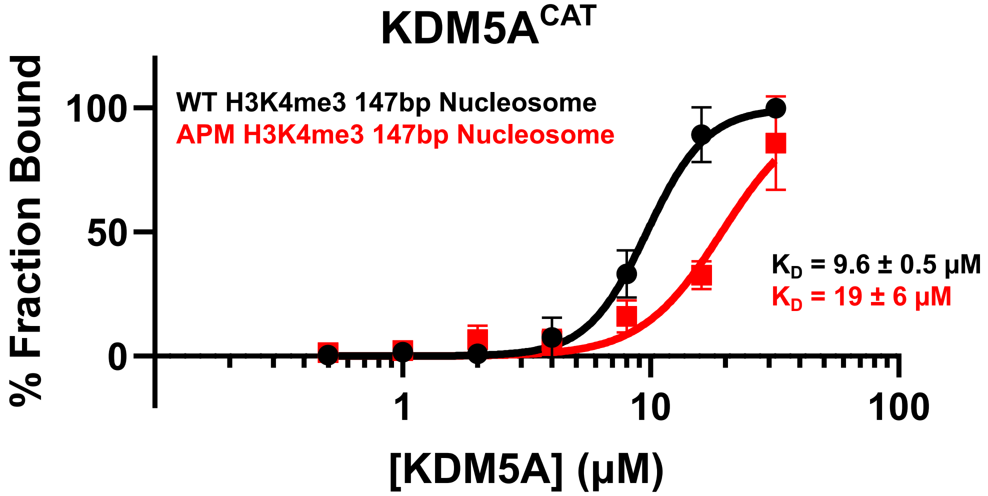


EMSA binding assay of wild-type KDM5A$^{CAT}$ to wild-type and acidic patch mutant H3K4me3 nucleosomes. Data are presented as the mean ± s.d. from three replicates collected across two independent experiments.

### Supplementary Figure S3

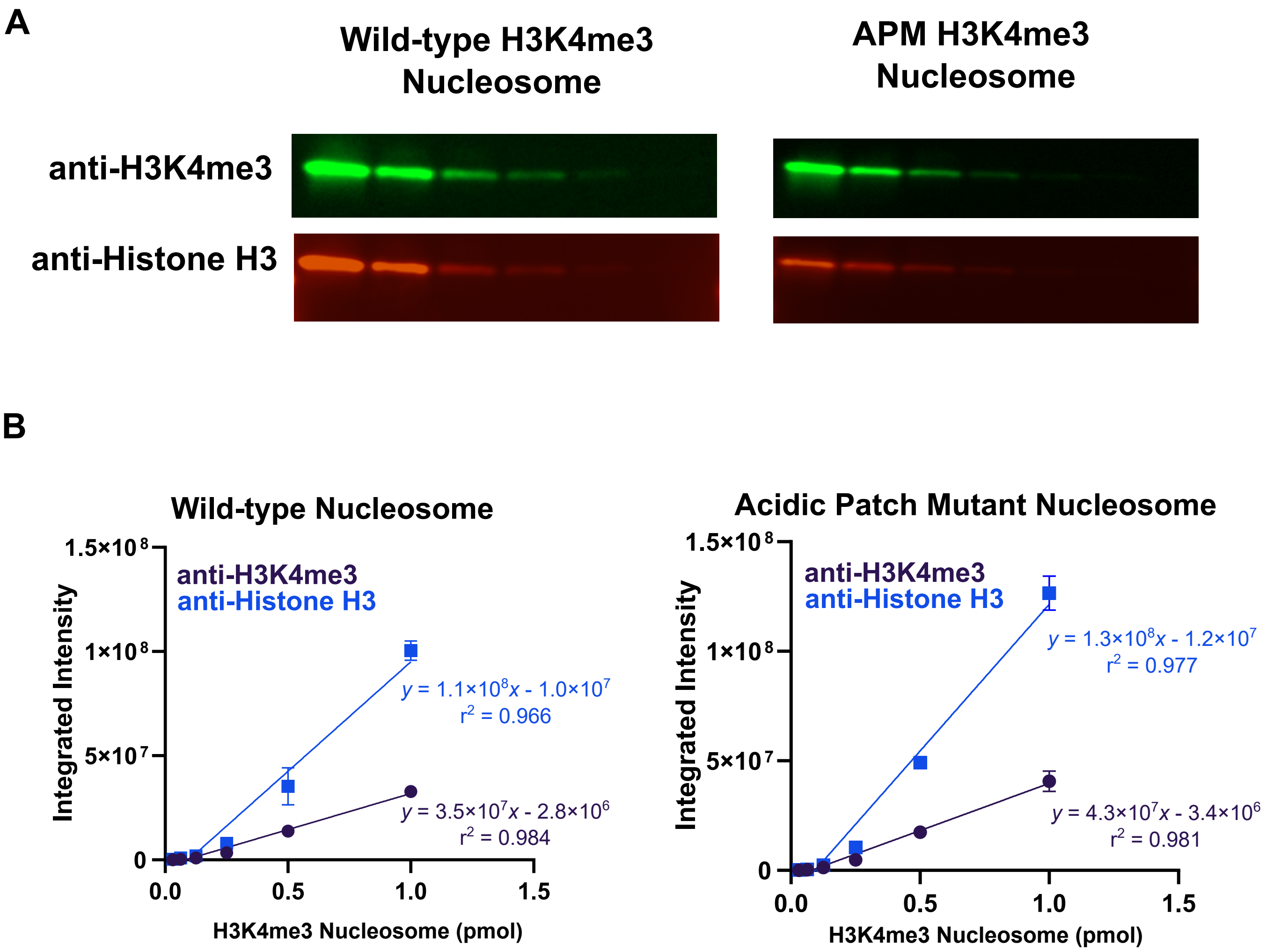


**A**. Western blot for quantification of the linear range of H3K4me3 and H3 antibodies. A serial dilution of wild-type and APM H3K4me3 nucleosomes ranging from 0-1 pmol was prepared.

**B.** Linear range of detection for H3K4me3 and H3 antibodies on wild-type and APM H3K4me3 nucleosomes. Antibody detection is linear within tested conditions (0 – 1 pmol, n=2). Nucleosome demethylation reactions were performed using 0.5 pmol of H3K4me3 nucleosome.

### Supplementary Figure S4

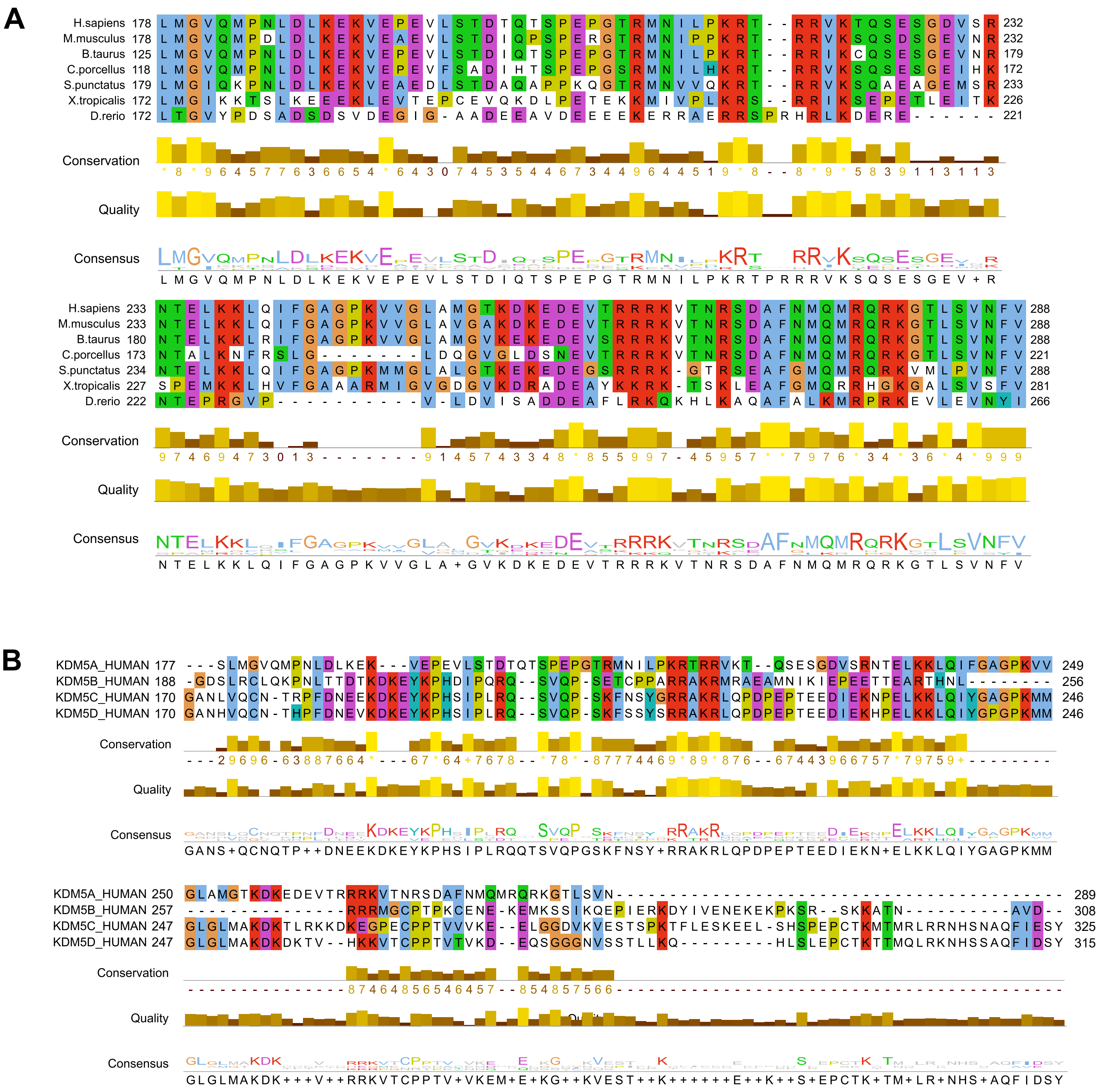

**A.** Multiple sequence alignment of KDM5A orthologs in vertebrates. Residues colored according to the ClustalX color scheme. Alignment performed using ClustalW

**B.** Multiple sequence alignment of the complete disordered regions of KDM5 family proteins. Residues colored according to the ClustalX color scheme. Alignment performed using ClustalW

### Supplementary Figure S5

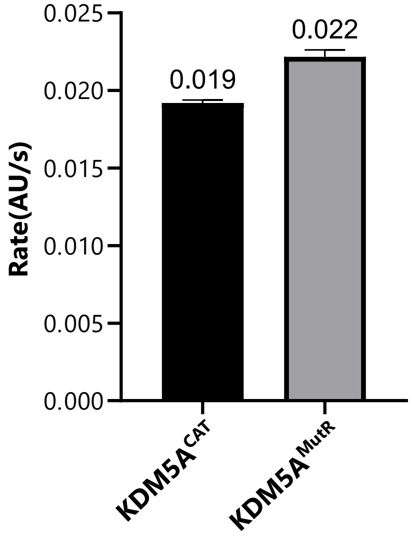


| | Rate (AU/s) |
|---|---|
| KDM5A$^{CAT}$ | 0.0192 ± 0.0002 |
| KDM5A$^{MutR}$ | 0.0221 ± 0.0004 |

Catalytic rates for demethylation of H3K4me3 peptides by KDM5A$^{CAT}$ and KDM5A$^{MutR}$. Demethylation kinetics were measured using a formaldehyde dehydrogenase assay, coupling the demethylation of H3 peptide substrate to the formation of formaldehyde. (n=3).

### Supplementary Figure S6

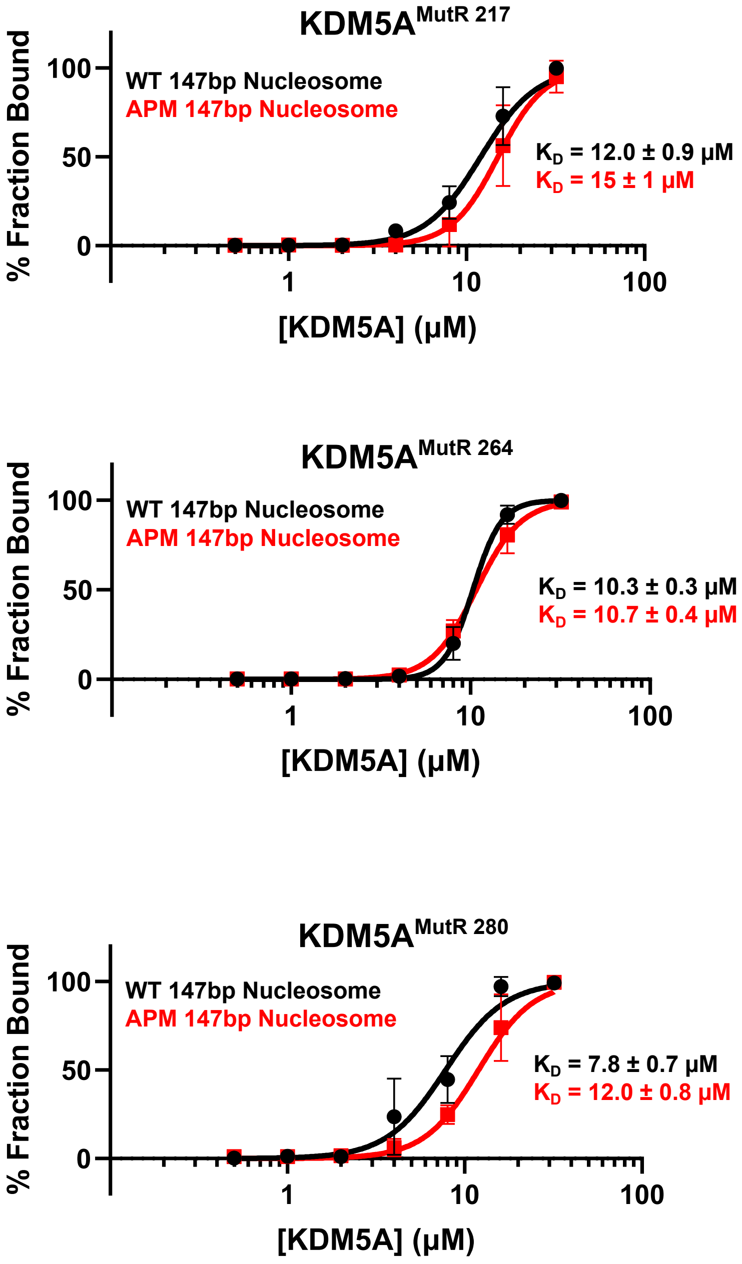


Quantification of EMSA binding assays of KDM5A arginine motif mutants to wild-type and acidic patch mutant nucleosomes presented in Figure 3B. For $KDM5A^{MutR\ 217}$ and $KDM5A^{MutR\ 264}$, data are presented as the mean ± s.d. from four replicates collected across two independent experiments. For $KDM5A^{MutR\ 280}$, data are presented as the mean ± s.d. from five replicates collected across three independent experiments.

### Supplementary Figure S7

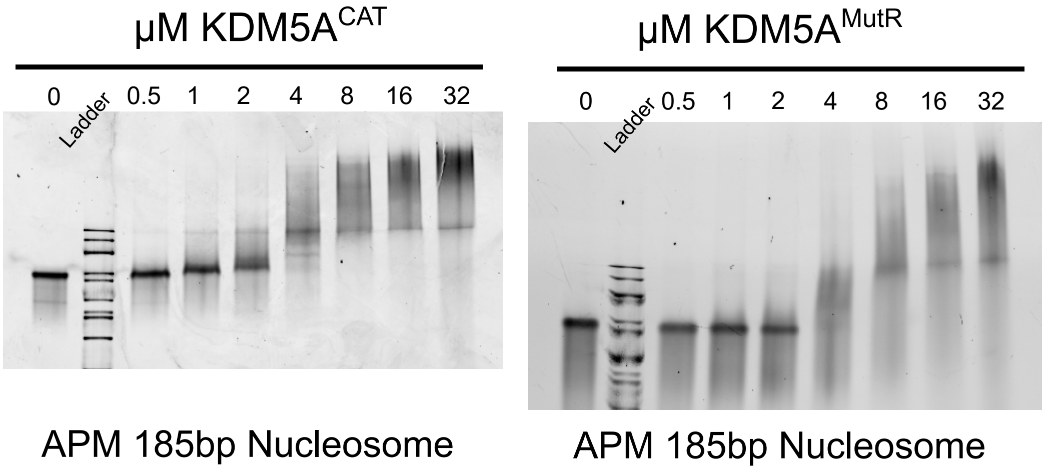


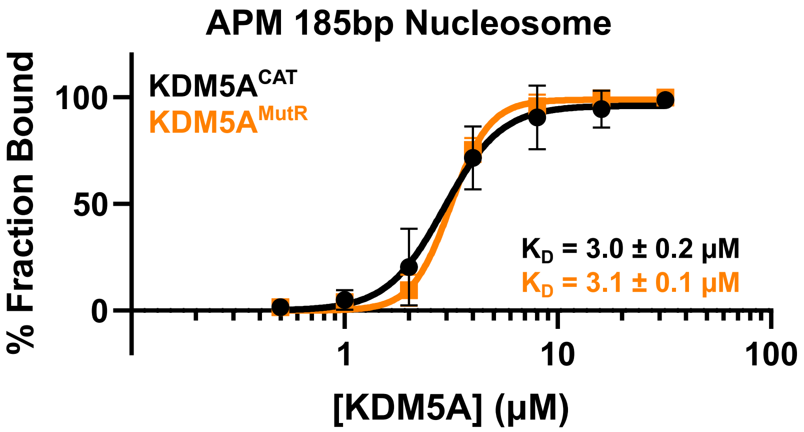


EMSA binding assay of wild-type $KDM5A^{CAT}$ and $KDM5A^{MutR}$ to acidic patch mutant 185bp nucleosomes. Data are presented as the mean ± s.d. from three replicates collected across two independent experiments.

#### Supplementary Figure S8

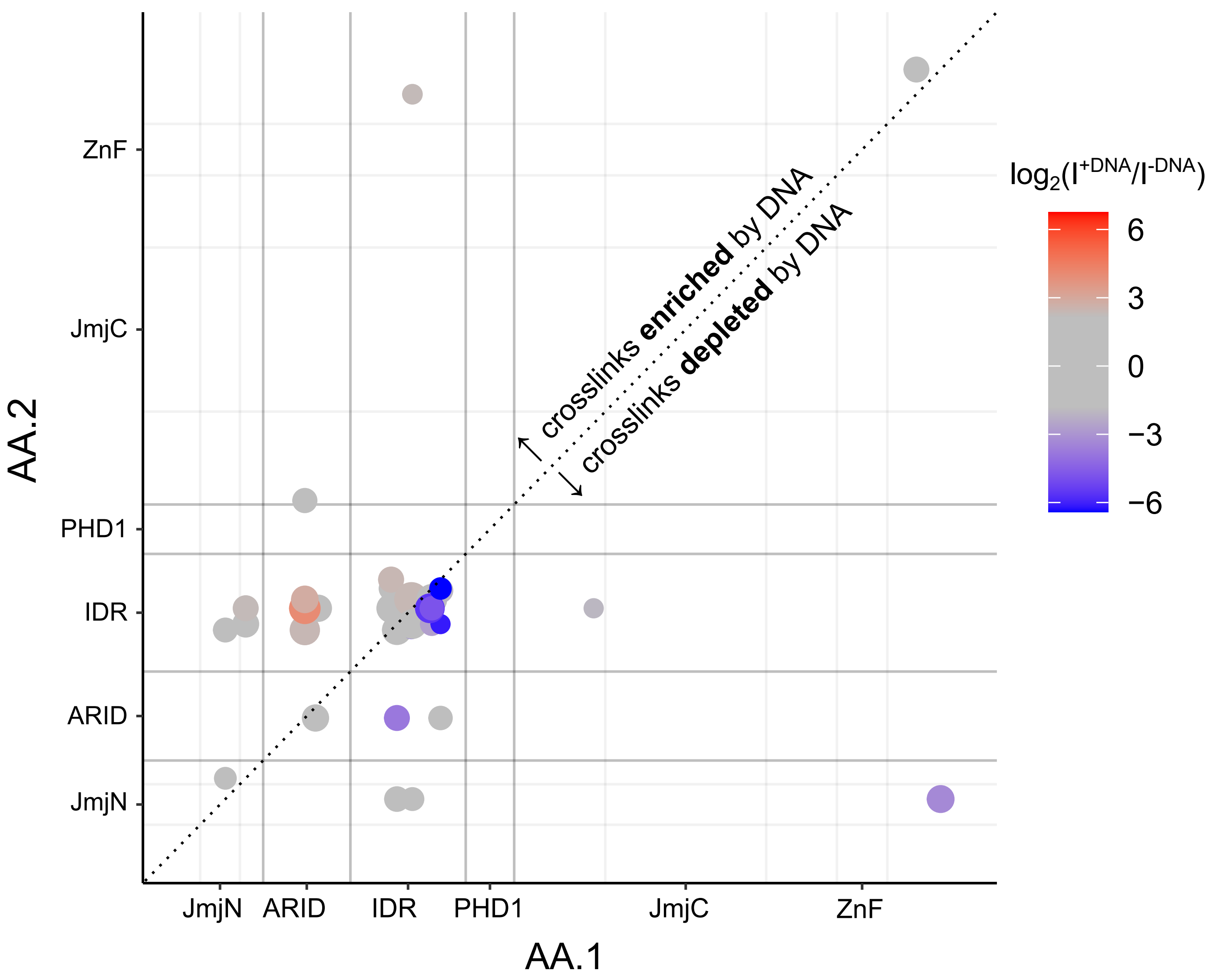


Two-dimensional map of quantified crosslinks in MS analysis of KDM5A$^{CAT}$, ±ARID-C1 dsDNA. Positions of the cross-link within KDM5A are given by the x and y-axes. KDM5A domain boundaries are demarcated by grey lines, with boundaries of the ARID and PHD1 domains emphasized. The size of points represents the relative intensity of the peptides as determined by MS, and points are colored based on the relative intensity with and without the presence of DNA. Enriched crosslinks in the presence of ARID-C1 dsDNA are positioned above the dividing line and colored red; depleted crosslinks are positioned below the dividing line and colored blue

### Supplementary Figure S9

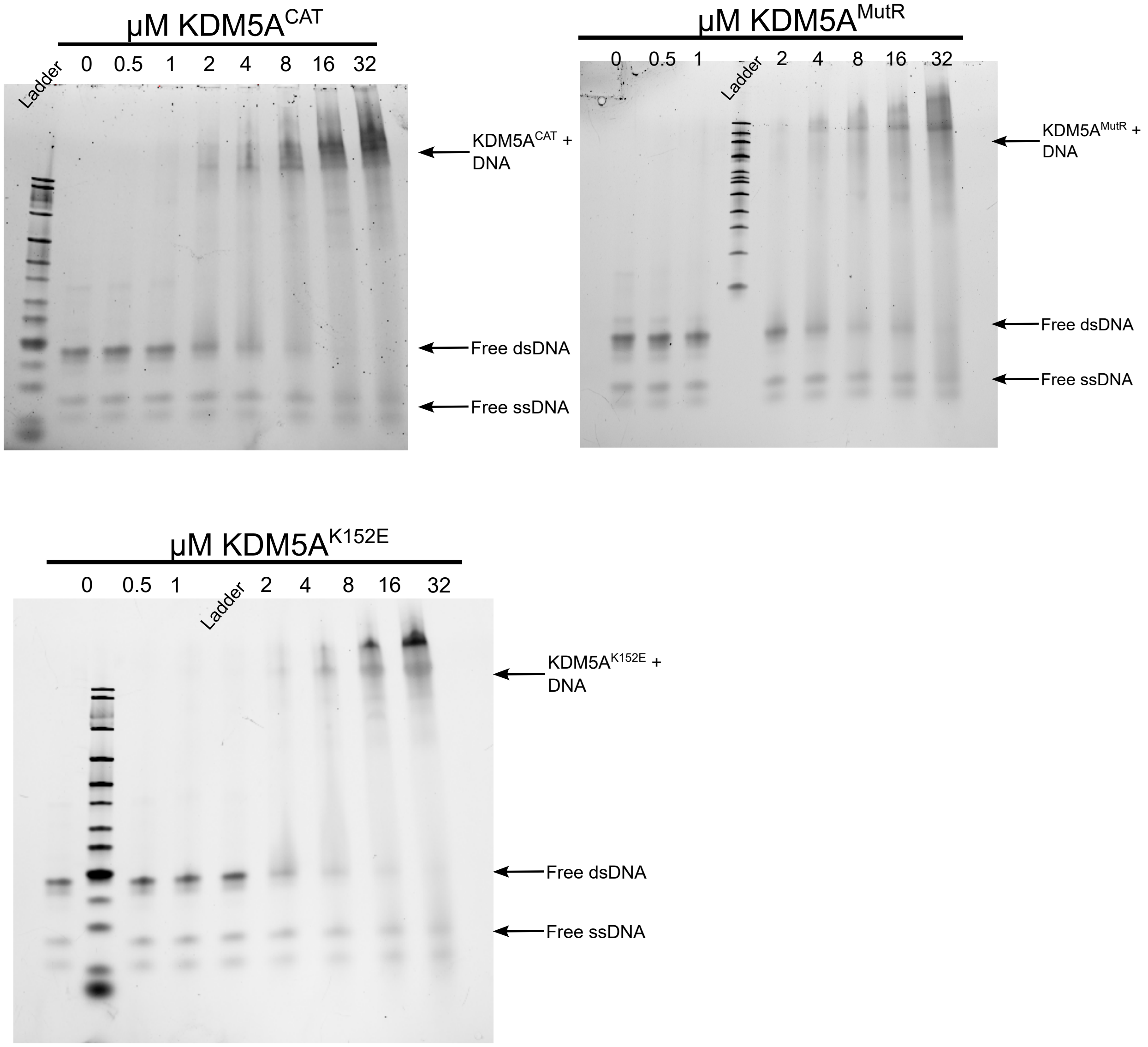


EMSA binding assay of wild-type $KDM5A^{CAT}$, $KDM5A^{MutR}$, and $KDM5A^{K152E}$ to double-stranded ARID-C1 DNA. Images for $KDM5A^{CAT}$ and $KDM5A^{K152E}$ are representative of three replicates collected across two independent experiments. The image for $KDM5A^{MutR}$ is representative of three replicates collected across two independent experiments.
